# A protease-initiated model of wound detection

**DOI:** 10.1101/2020.12.08.415554

**Authors:** James T. O’Connor, Aaron C. Stevens, Erica K. Shannon, Fabiha Bushra Akbar, Kimberly S. LaFever, Neil Narayanan, M. Shane Hutson, Andrea Page-McCaw

**Affiliations:** Department of Cell and Developmental Biology, Vanderbilt University, Nashville, Tennessee; Program in Developmental Biology, Vanderbilt University, Nashville, Tennessee; Department of Physics and Astronomy, Vanderbilt University, Nashville, Tennessee; Department of Biological Sciences, Vanderbilt University, Nashville, Tennessee; Institute for Integrative Biosystems Research and Education, Vanderbilt University, Nashville, Tennessee; Vanderbilt-Ingram Cancer Center, Vanderbilt University, Nashville, Tennessee

## Abstract

Wounds trigger surrounding cells to initiate repair, but it is unclear how cells detect wounds. The first known wound response of epithelial cells is a dramatic increase in cytosolic calcium, which occurs within seconds, but it is not known what initiates this calcium response. Specifically, is there an instructive signal detected by cells surrounding wounds? Here we identify a signal transduction pathway in epithelial cells initiated by the G-protein coupled receptor Methuselah-like 10 (Mthl10) activated around wounds by its cytokine ligands, Growth-blocking peptides (Gbps). Gbps are present in unwounded tissue in latent form, requiring proteolytic activation for signaling. Multiple protease families can activate Gbps, suggesting it acts as a detector to signal the presence of several proteases. We present experimental and computational evidence that proteases released during cell lysis serve as the instructive signal from wounds, liberating Gbp ligands to diffuse to the Mthl10 receptors on epithelial cells and activate downstream release of calcium. Thus, the presence of a nearby wound is signaled by the activation of a Gbp protease detector, sensitive to multiple proteases released after cellular damage.

## Introduction

When tissue is wounded, the surrounding cells rapidly respond to repair the damage. Despite the non-specific nature of cellular damage, there is remarkable specificity to the earliest cellular response: cells around the wound increase cytosolic calcium, and this damage response is conserved across the animal kingdom^1–12^. The calcium response is not limited to epithelial cells at the wound margin, but extends even to distal cells^1,2,4,5^. Additionally, multiple molecular mechanisms downstream of calcium have been identified that regulate wound-induced gene expression or cell behavior^1,2,4,5,13–16^. However, it remains unclear how the calcium response is initiated, i.e., how cells detect wounds. Here we investigate the molecular mechanisms by which the wound initiates cytosolic calcium signals that in turn initiate repair or defense responses.

Calcium has been well-established as a versatile and universal intracellular signal that plays a role in the modulation of numerous intracellular processes^13,14,17,18^ including cell migration and proliferation^7,19^. In fact, calcium is a key regulator of actomyosin dynamics^1,13,20–23^ and cell proliferation^19,24,25^ and is implicated in JNK pathway activation^15^, which is required for proper wound repair^26–30^. Thus, the immediate increase in cellular calcium may promote many aspects of wound repair.

Although an increase in cytosolic calcium is necessary for wound repair^5–7,13,14^, there is less clarity on the mechanisms that trigger the increased cytosolic calcium in cells near to and distant from the wound. Some studies have shown that wound-induced cytoplasmic calcium comes from the extracellular environment^1,2,7,8,11,13–15^, entering the cell either directly upon plasma membrane damage or through calcium ion channels, of which TRPM has been of particular interest. Others have shown that calcium is released from the endoplasmic reticulum (ER), mediated by the IP_3_ Receptor^6,7,9,12,31,32^, initiated by an unknown G-Protein Coupled Receptor (GPCR) or Receptor Tyrosine Kinase (RTK). Others still have indicated that wound-induced calcium responses are initiated by the mechanical stimulus of wounding^32–35^. Elucidating the mechanisms by which calcium signaling is triggered *in vivo* is critical to understanding how wound information is transmitted through a tissue in order to change cellular behavior and repair the wound.

Using an *in vivo* model, we previously showed that damaged cells within wounds become flooded within milliseconds by extracellular calcium entering through microtears in the plasma membrane^8^. Although this calcium influx expands one or two cell diameters through gap junctions, it does not extend to more distal cells. Strikingly, after a delay of 45-75 seconds, a second independent calcium response expands outward to reach a larger number of distal cells. Here we identify the relevant signal transduction pathway initiated by activation of the GPCR Mthl10. Downstream, signals are relayed through Gaq and PLCβ to release calcium from the ER. Upstream, Mthl10 is activated around wounds by the cytokine ligands Growth Blocking Peptides (Gbps). Further, we provide experimental and computational evidence that the initiating event for the distal calcium response *in vivo* is a wound-induced release of proteases that activate the latent Gbp cytokines, cleaving them from inactive/pro-forms into cleaved/active forms.

## Results and Discussion

To investigate the calcium responses after wounding, we analyzed *Drosophila* pupae (Fig. 1A), which are amenable to live imaging *in vivo* because they are stationary throughout development. By partially removing the pupal case, we were able to access and wound the epithelial monolayer of the notum, or dorsal thorax (Fig 1A, white box). After wounding and imaging, nearly all wild-type pupae recovered to eclose as adults.

**Figure 1.**
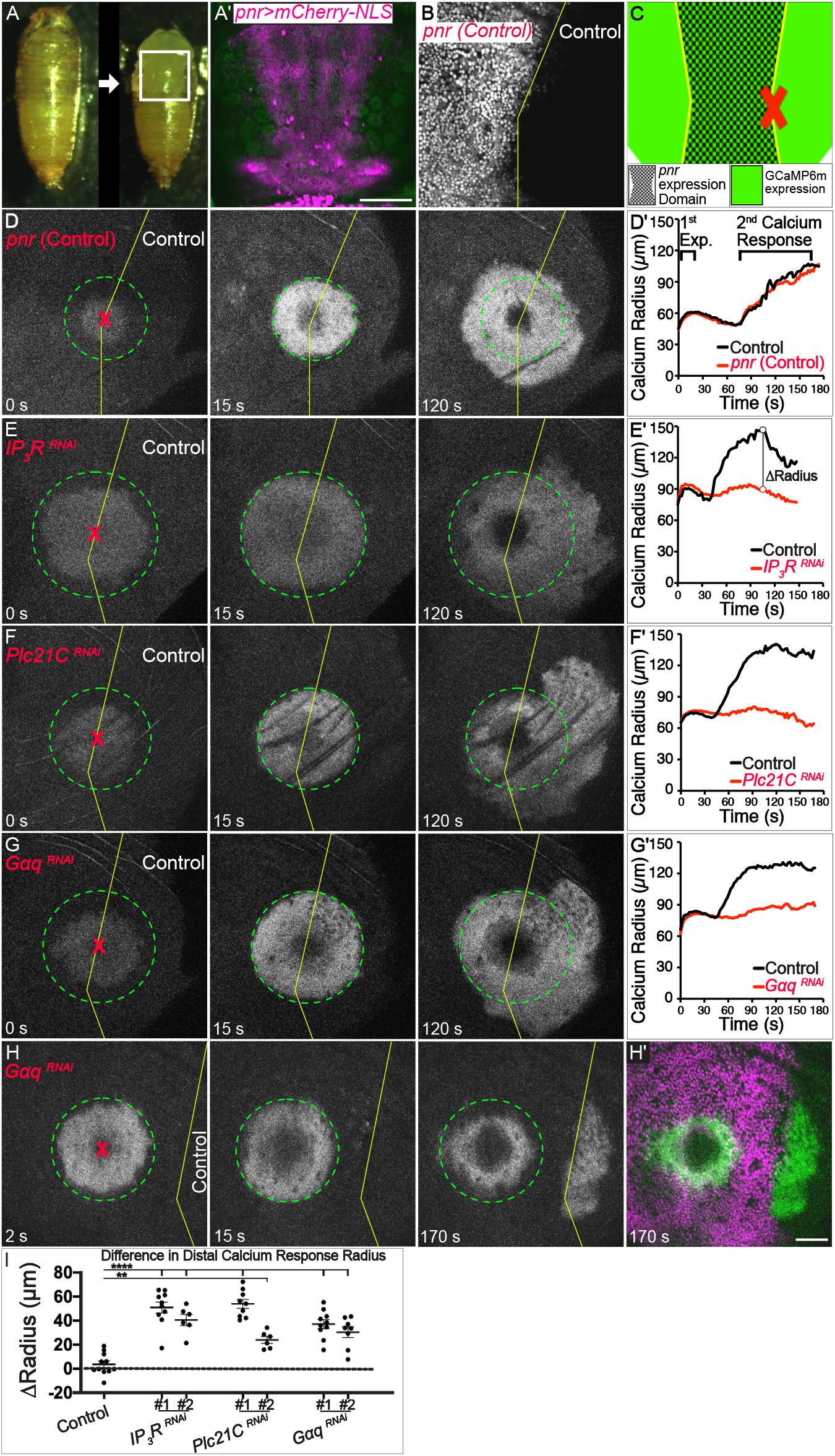
Wounds trigger calcium release via the Gq pathway. **(A)** Experimental model, *Drosophila* pupa (left) with pupal case removed (right). Gene expression is controlled in the *pnr* domain (magenta, **A’**). **(B-C)** wounds (red X) are targeted at the *pnr* boundary (yellow line, **B, C**). (**D-H)** GCaMP6m calcium reporter. **(D, D’)** Calcium response to wounds in the absence of gene knock-down. The max radius of the rapid first calcium expansion is marked by green circle; the distal calcium response begins ~45-75 s after wounding. Analysis of the calcium radius with respect to time **(D’)** demonstrates that the response is symmetric on both sides of the *pnr* boundary. (**E–G**) The distal calcium response requires Gq-pathway components knocked down on the left side (*pnr* domain) of each panel. (**D’–G’**) Quantification of calcium dynamics for control (black) and *pnr* (red) sides of wounds. (**H**) Wound targeted within the *G_aq_* knock-down domain (left of yellow line) yields no distal calcium response until the initiating signal reaches the nearby control domain. (**H’**) Green shows calcium (*GCaMP6m*), magenta shows *pnr* domain where *G_aq_* is knocked down. (**I**) ΔRadius, difference in calcium signal radii at maximum extent of the distal calcium response for control minus *pnr* side of each wound (as shown in E’), with *pnr*-knockdown genotypes indicated; bars = mean ± SEM. Statistical analysis by one-way ANOVA, multiple comparisons WRT control group, **p<0.01, ****p<0.0001. Scale bars = 200 μm (A’), or 50 μm (B–H’).

The calcium responses observed after wounding are temporally and spatially complex and somewhat variable^8^, making it more difficult to identify underlying mechanisms by comparing wild-type and mutant animals. To circumvent this problem, we exploited the radially symmetry of the calcium response by manipulating gene expression in only one part of the wound, allowing us to assess symmetry in control vs. experimental regions. We used the *Gal4/UAS* system to manipulate gene expression in the central region of the notum with *pnr-Gal4* (Fig. 1A’), and then wounded on the margin between the control and knock-down region, so that half the wound served as an internal control (Fig. 1B, C). We monitored the symmetry of the calcium responses with a ubiquitously expressed calcium reporter, *actinP-GCaMP6m,* comparing the experimental (*pnr*) domain with the adjacent internal control. When no other genes are manipulated, the calcium signals remain radially symmetric about the wound (Fig. 1D, Movie 1).

### The distal calcium response signals to release calcium via IP_3_

We first used this internally controlled system to knock down gap junctions, which we have previously shown are required for the first calcium expansion^8^. As expected, the first expansion did not occur in the gap junction knockdown region but did in the control region (Fig. S1A). Importantly, the second distal calcium response still occurred in the gap junction knockdown region, although it appeared “speckled” because each cell transduced the signal independently of its neighbors (Fig. S1A), as has been reported previously^8,32,36^. This result indicated that the initiation of the distal calcium response requires neither gap junctions nor the first (gap junction-dependent) expansion.

To determine whether the calcium of the distal response was coming from internal stores within the cell, we knocked down IP_3_ Receptor (IP_3_R), which controls calcium release from the endoplasmic reticulum. IP_3_R was absolutely required for the distal calcium response (Fig. 1E). Importantly, the effect was limited to the experimental domain, while the control domain calcium response remained intact. We confirmed this result with two independent RNAi lines and an IP_3_ sponge (Fig. S1B) and concluded that calcium is released from intracellular stores in an IP_3_ dependent manner. Based on this result and previous literature^6,37^, we tested the role of G_q_-signaling by knocking down PLCβ or G_αq_, and found that each was required for the distal calcium response (Fig. 1F, G, Movie 2). Like laser wounds, puncture wounds also had two distinct calcium waves: the first occurred immediately and required gap junctions, whereas the second was delayed by tens of seconds and required the G_q_ pathway (Movie 3). Thus, the mechanisms underlying the two calcium responses appear to be general wound responses, not specific to laser ablation.

Interestingly, a modification of our internally controlled system suggested that wound-induced signals can diffuse in the extracellular space. Rather than wounding on the *pnr* domain border, we wounded in the middle of the *pnr* domain where G_aq_ was knocked down. As expected, no distal calcium response occurred within the knockdown domain, but the calcium response did jump the gap to suddenly appear in the even more distal control domain (Fig. 1H, Movie 4). To quantify how gene knockdown alters the distal calcium response, we subtracted the maximum calcium radius of the experimental region from the radius of the control region (illustrated in 1E’). Each Gq-pathway knockdown significantly inhibited the calcium response (Fig. 1I). Taken together, the distal calcium response occurs through a Gq-signaling pathway likely activated by a diffusible signal.

### The distal calcium response requires the GPCR Mthl10

Before screening potential G-protein coupled receptors that may be activating the Gq-pathway, we used our internally-controlled system to test the TRPM ion channel, previously reported to alter the wound-induced calcium response in this pupal tissue^1,2,6^. Knockdown of TRPM with either of the two functional RNAi lines previously used gave no discernible effect on the calcium response: the experimental domain remained symmetrical to its internal control (Fig. 2A, D). Whereas previous study focused on GCaMP intensity, our analytical framework is optimized to identify spatial or temporal changes in the calcium response to wounds, which may explain the difference in our results.

**Figure 2:**
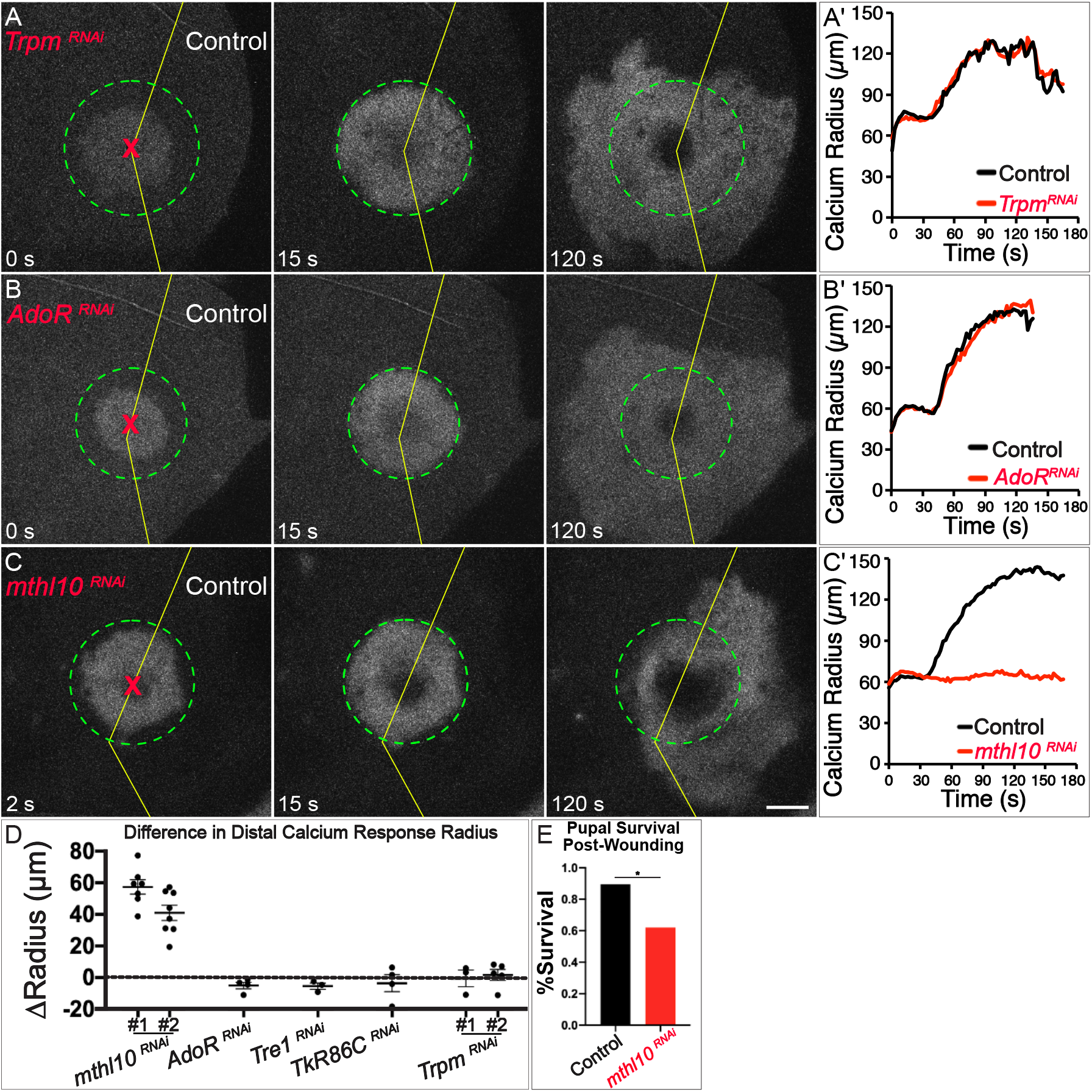
The distal calcium response requires the GPCR Methuselah-like 10 (Mthl10) (**A–C**) GCaMP6m reporter showing cytosolic calcium in representative samples of various knockdowns in the *Drosophila* notum split-expression system. Gene manipulations are performed in the domain of *pnr* expression (left side) and wounds (red X) are administered at the *pnr* domain boundary (yellow line). Maximum extent of first expansion is marked by dashed green circle. (**A**) Knockdown of the calcium ion channel *TRPM* has no effect on either calcium response. (**B**) Knockdown of the GPCR Adenosine Receptor *(AdoR)* has no effect on either calcium response. (**C**) Knockdown of the GPCR *Methuselah-like 10 (Mthl10)* phenocopies the Gq-pathway knockdowns, with a dramatic decrease in the distal calcium response. (**A’–C’**) Quantification of calcium signal radius versus time in on control (black) and *pnr*-expression (red) sides of each wound. (**D**) ΔRadius, with *pnr*-knockdown genotypes indicated; bars = mean ± SEM. (**E**) Pupal survival past eclosion after large wounds. Quantification by Chi-Square, n=29 each, p=0.0141. Scale bars = 50 μm.

The requirement of Gq suggested the distal calcium response is initiated by a GPCR. To identify the GPCR controlling the distal calcium response, we knocked down candidates, prioritizing receptors with known epithelial expression and Gq activity^38^. Although cell-culture studies implicated ATP as a ligand activating calcium after wounding^37^, and a recent study in mice implicated ADP in intercellular calcium waves following viral infection^39^, knockdown of *AdoR,* the only *Drosophila* adenosine receptor, had no effect on the calcium response (Fig. 2B,D). Additionally, knockdown of GPCRs *Tre1* or *TkR86C,* both implicated in wound responses^40,41^, had no effect on the calcium response (Fig. 2D). Remarkably, knockdown of the cytokine-activated GPCR *Methuselah-like 10 (Mthl10)^42^* completely eliminated the distal calcium response (Fig. 2C,D, Movie 5). We confirmed this result with two independent RNAi lines. To test whether *Mthl10* is important for wound recovery, we measured pupal survival after large wounds and found that *Mthl10* knockdown throughout the notum epithelium decreased pupal survival by ~30% (Fig. 2E).

### Mthl10 is activated by Gbp1 and Gbp2

Mthl10 is known to be activated by a class of cytokines known as Growth-blocking peptides (Gbps), of which there are five in *Drosophila*^42,43^. Gbps are secreted by the Drosophila fat body in a latent pro-peptide form, requiring proteolytic cleavage for activation^43–47^. Because Gbps are secreted and thus non cell-autonomous, the internally controlled knockdown approach was not effective at testing their role (data not shown); thus, we tested flies homozygous for *Df(2R)Gbp-ex67,* a small chromosomal deletion removing the neighboring genes *Gbp1* and *Gbp2* ^48^ (hereafter referred to as *ΔGbp1,2;* Fig. 3A). Upon wounding, *ΔGbp1,2* pupae lacked the distal calcium response (13 out of 14 pupae, Fig. 3B). To test the roles of these Gbps individually, we used CRISPR to generate individual null mutants of *Gbp1* or *Gbp2* (Fig. 3A). Each mutant displayed the distal calcium response (Fig. 3C, D), indicating that each can signal independently and they function redundantly. Thus, Gbp1 and Gbp2 are both extracellular ligands that relay wound information to surrounding cells.

**Figure 3:**
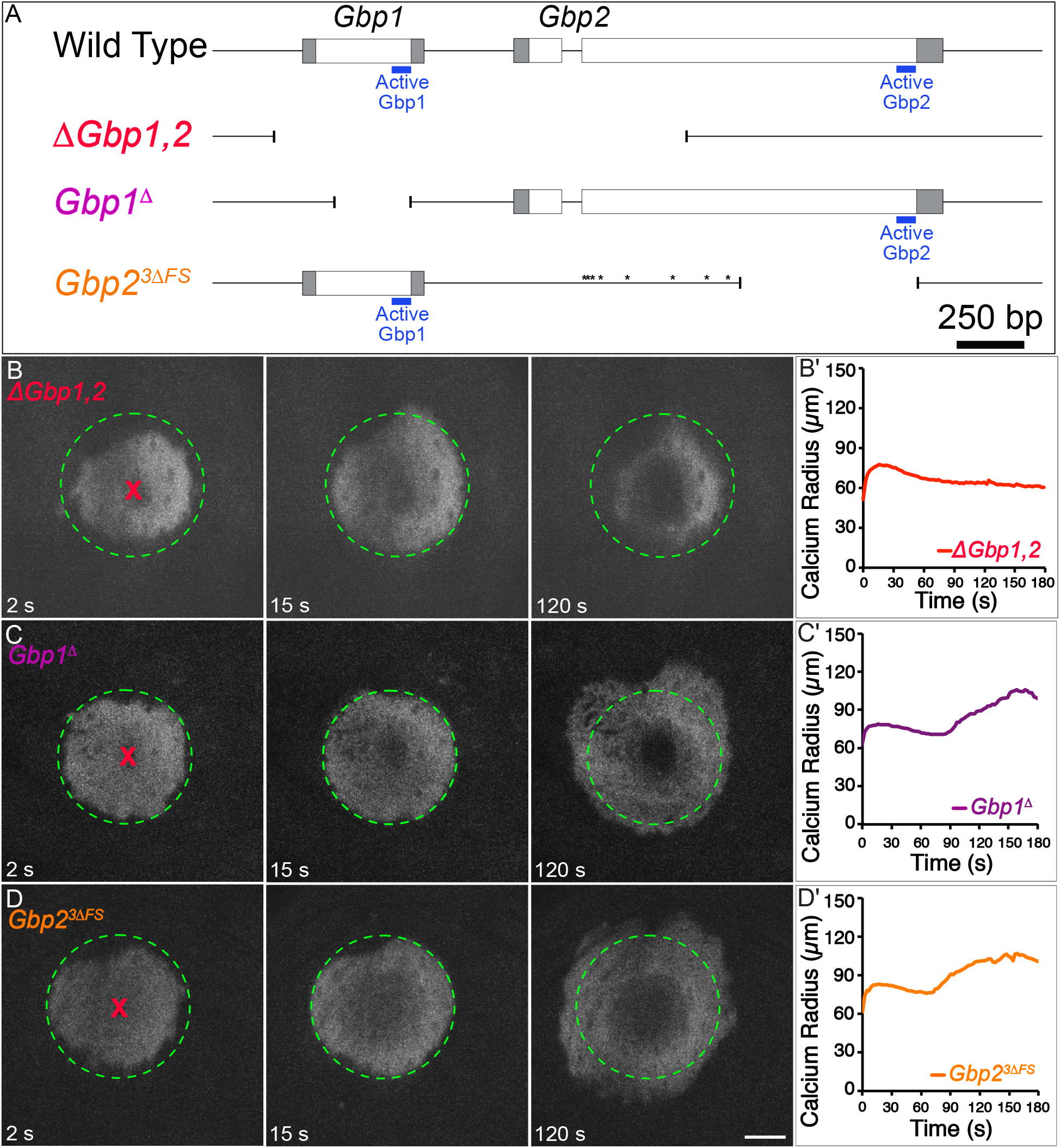
The distal calcium response requires either Gbp1 or Gbp2. (**A**) Schematic showing the genomic region for *Gbp1* and *Gbp2,* the deletion *ΔGbp1,2*, and the *Gbp1* or *Gbp2* null mutants generated for this study (transcription proceeds from left to right). The *Gbp1^Δ^* deletion allele is missing 282 bases within the coding region of *Gbp1,* removing most of the region encoding the Gbp1 active peptide (blue). The complex *Gbp2^3ΔFS^* allele comprises 8 point mutations (asterisks) and 3 deletions, inducing frameshifts and premature terminations upstream of the Gbp2 active peptide (blue) (see Methods). (**B–D**) GCaMP6m reporter showing cytosolic calcium in representative samples of various knockouts in the *Drosophila* notum. (**B, B’**) The distal calcium response is absent after wounding homozygous *ΔGbp1,2.* n=13/14. (**C-D**) Homozygous *Gbp1^Δ^* (**C**) or *Gbp2^3ΔFS^* (**D**) each retain the distal calcium response (n = 13 and 7 respectively). Scale bar = 50 μm.

If Gbps are wound-induced signals that direct cells to increase calcium, then ectopic Gbp would be expected to activate a calcium response, even without a wound. Unfortunately, the pupal notum is protected by an impermeable cuticle that makes it impossible to add Gbp directly to the tissue, and it cannot be removed without wounding. We thus turned to larval wing discs, sacs of epithelial tissue that have no cuticle. We applied synthetic active-form Gbp peptides directly to wing discs *ex vivo* at varying concentrations. Excitingly, we found that both Gbp1 and Gbp2 elicited a strong calcium response in wing discs when added *ex vivo* at 5 nM or 50 nM, respectively (Fig. 4A, B). Gbp1 and Gbp2 did not act synergistically: when added together, they were no more potent than when added individually (Fig. S2A). To test whether these responses were Mthl10-dependent, we mounted one control and one *Mthl10* knockdown disc side-by-side in a media bubble and added Gbp peptide to both discs simultaneously. As expected, the calcium responses were absent from the *Mthl10* knockdown discs (Fig. 4C, D, Movie 6).

**Figure 4:**
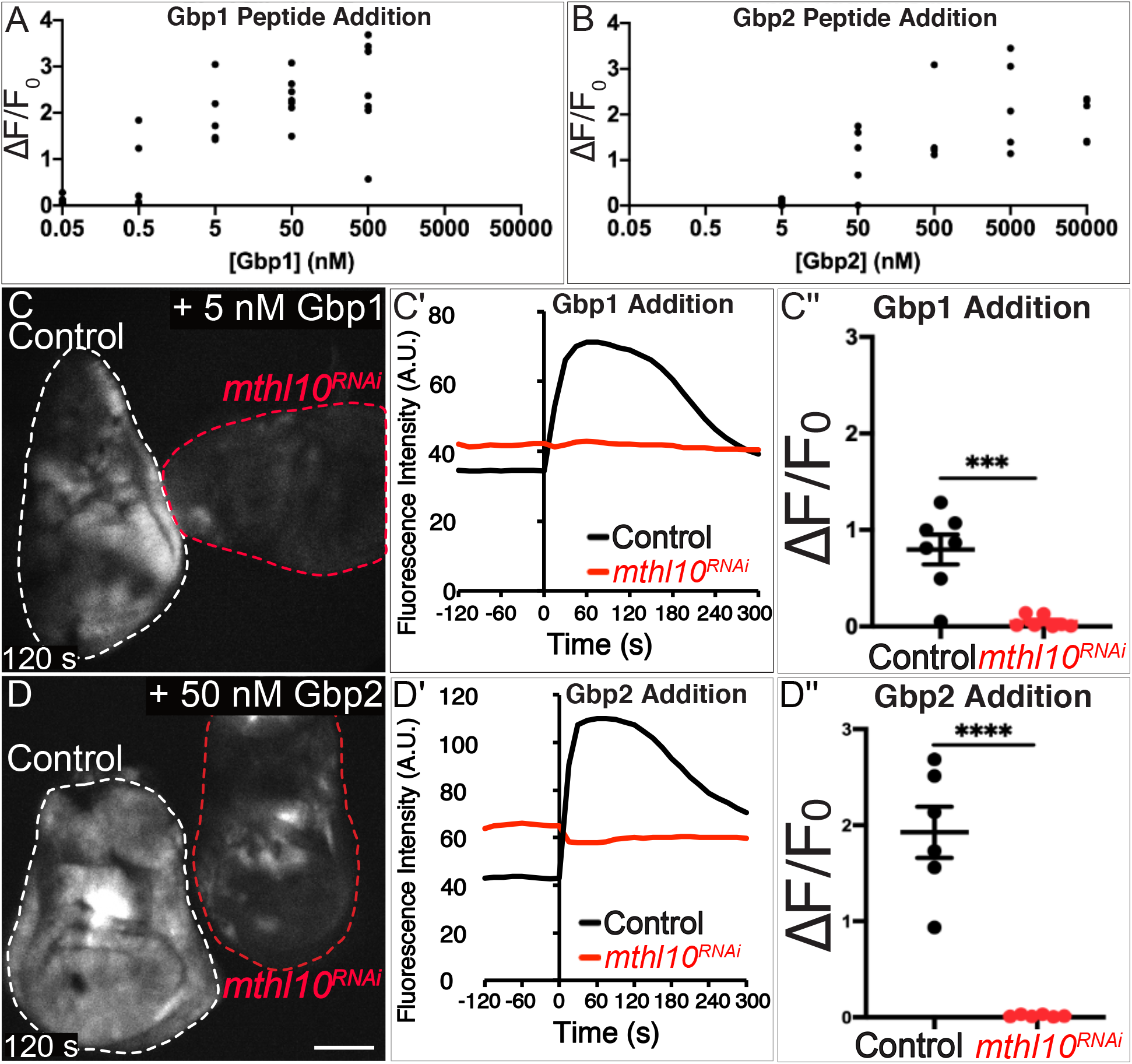
Gbp1 and Gbp2 peptides elicit calcium responses in wing discs through Mthl10. (**A-B**) Gbp1 or Gbp2 peptide elicit a concentration-dependent calcium response in *Drosophila* wing discs. 5 nM Gbp1 or 50 nM Gbp2 consistently elicits a calcium response. (**C-D**) Assays were performed with two wing discs (one control and one *mthl10* knockdown) mounted adjacently and activated simultaneously. *Mthl10* is required in wing discs for Gbp1 (**C**) or Gbp2 (**D**) to activate a calcium response. Change in GCaMP6m fluorescence over time for samples **C** and **D** is quantified in **C’** and **D’**, respectively. Normalized change in fluorescence is quantified for all samples in **C’’** and **D’’**. Scale bar = 100 μm. Graph bars represent mean and SEM. ***p<0.001, ****p<0.0001, by Student’s t-test.

Finally, we asked whether the three other Gbps could elicit calcium in wing discs and found that Gbp4 and Gbp5 could elicit calcium consistently at 50 nM, (and Gbp4 occasionally even at 5 nM), while Gbp3 was hardly effective, even at the maximum concentration tested of 50 μM (Fig. S2B). Despite the activity of these other Gbps *ex vivo,* the loss of the distal calcium response in the *ΔGbp1,2* pupae indicates Gbp1 and Gbp2 are responsible for the calcium increase in the pupal notum.

### Gbps and Mthl10 are required for calcium waves in wing discs

Previous studies have shown that wing discs cultured *ex vivo* displayed potent calcium responses upon exposure to fly extract^32,35,36^. Because fly extract is created by lysing and homogenizing whole flies, we hypothesized that it contained wound-induced signals that activate wound-detection pathways in wing discs to initiate calcium responses. Indeed, we observed the calcium response was *mthl10* dependent by adding fly extract to control and *mthl10* knockdown wing discs simultaneously (Fig. 5A, C, Movie 7). Since extract of adult flies is predicted to contain both Gbp4 and Gbp5, which may be confounding variables, we also tested extract made from wild-type larvae, a stage that expresses only Gpb1-3^43^; like adult extract, larval extract activated calcium responses in wing discs in an *mthl10*-dependent manner (Fig. 5B,C, Movie 7).

**Figure 5:**
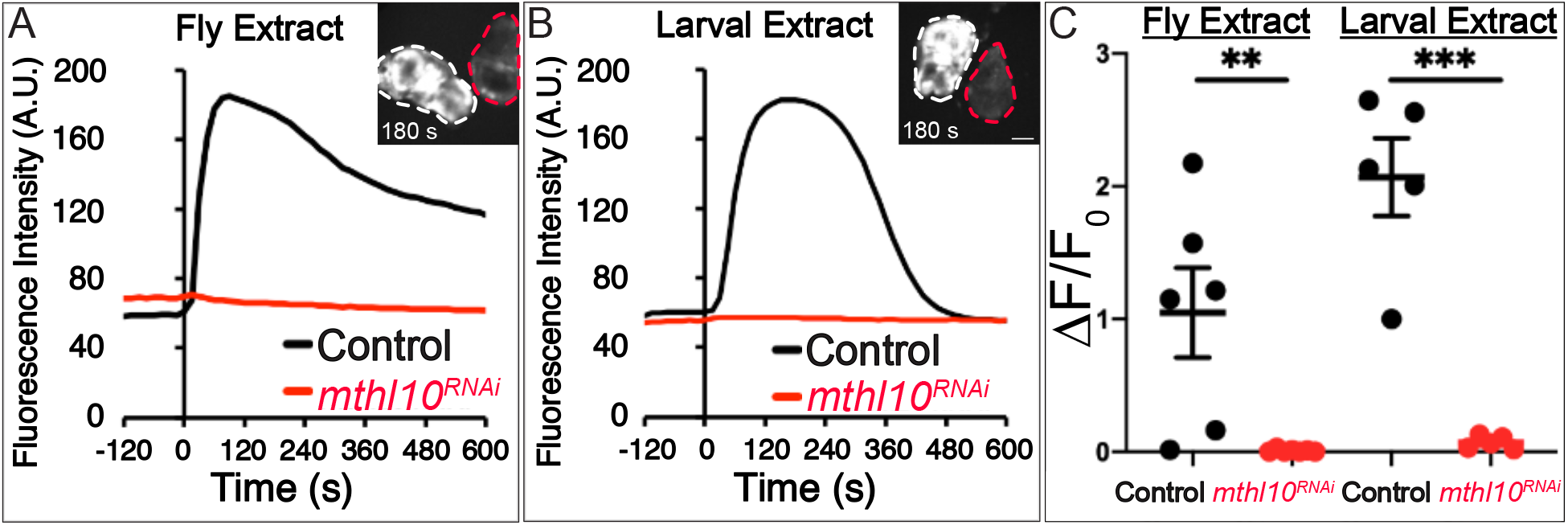
*Drosophila* extracts elicit a calcium response in wing discs through Mthl10. (**A-B**) Assays were performed with two wing discs (one control and one *mthl10* knockdown) mounted adjacently and activated simultaneously. *Mthl10* is required in wing discs for 5% fly extract (**A**) or 5% larval extract (**B**) to activate a calcium response. Normalized change in fluorescence is quantified for all samples in **C**. Scale bar = 100 μm. Graph bars represent mean and SEM. **p<0.01, ***p<0.001, by Student’s t-test.

To determine whether the calcium-activating signal in the extract is Gbp, we added extract made from homozygous *ΔGbp1,2* larvae to homozygous *ΔGpb1,2* wing discs. We found that no calcium response occurs when Gbps are absent from both disc and extract (Fig. 6A, C, Movie 8). This result demonstrates that the calcium response to larval extract requires Gbp1 and/or Gbp2. To determine whether Gbp is supplied by the extract itself or by the disc, we added wildtype extract to *ΔGbp1,2* mutant discs and mutant extract to wild-type discs. Surprisingly, both were able to activate calcium signaling (Fig. 6A-C, Movie 8). The former result is expected: extract made from wild-type larvae contains Gbp, which elicits calcium when applied to wing discs. The latter result is less intuitive: it suggests that the wing discs themselves maintain a local supply of Gbp that can be activated by larval extract. If the discs already have Gbp, then what does the extract provide to activate calcium? This result suggested that proteases in the extract are necessary to cleave latent pro-Gbp from the disc to elicit the calcium response.

**Figure 6:**
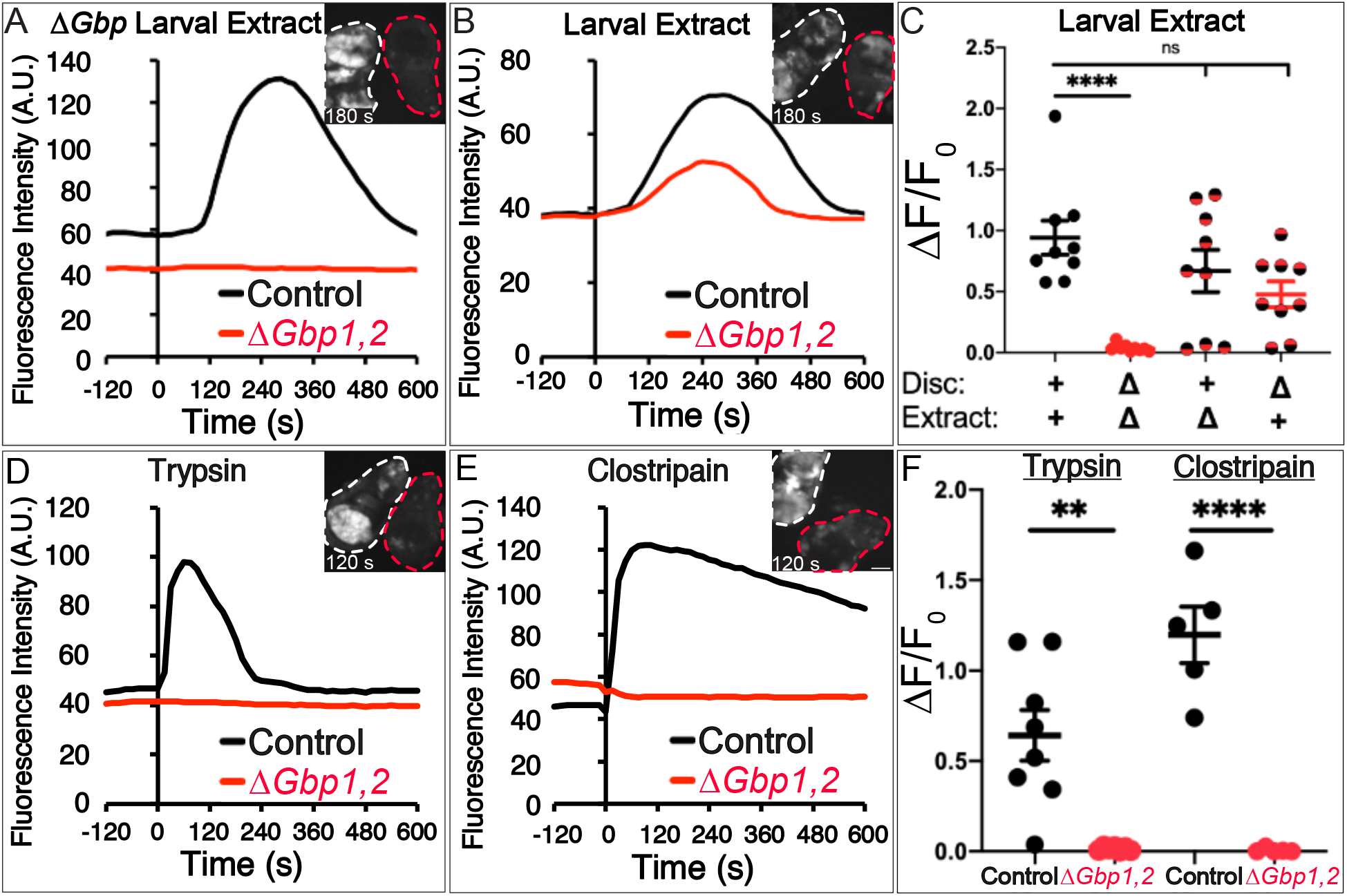
*Drosophila* extracts elicit calcium responses through Gbp and are activated by proteases. (**A-F**) All assays were performed with two wing discs (one control and one *ΔGbp1,2)* mounted adjacently and activated simultaneously (A, B, D, E, insets). (**A-C**) Gbp is required for the larval extract-mediated calcium response, as no response occurs if Gbp is absent from both extract and wing disc (**A**). The signal can be supplied by either the wing disc (**A**) or by the larval extract (**B**). Normalized change in fluorescence is quantified in **C**. **(D-F)** Serine protease trypsin (**D**) or cysteine protease clostripain (**E**) activate calcium in a Gbp-dependent manner. Normalized change in fluorescence is quantified in **F**. Scale bar = 100 μm. Graph bars represent mean and SEM. **p<0.01, ****p<0.0001 by two-way ANOVA in **C** and Student’s t-test in **F**.

### Gbps are activated by multiple proteases

Bioinformatics analysis predicted that both pro-Gbp1 and pro-Gbp2 could release their C-terminal active cytokine peptides after cleavage by several unrelated proteases. To confirm the activity of proteases in the lysate, we tried to inhibit them using broad-spectrum protease inhibitors, singly and in combination. Unfortunately, in most cases the vehicle (DMSO) or the inhibitor alone induced calcium waves in wing discs, stymieing our ability to use inhibitors in this experiment. As an alternative approach, we noted that the serine protease trypsin and the cysteine protease clostripain are both predicted to cleave Gbp1^49^. To test their sufficiency to activate calcium in wing discs, we added trypsin to wing discs and found that it elicited a calcium response in control but not ΔGbp1,2 wing discs (Fig. 6D, F). Similarly, adding clostripain elicited a calcium response in control but not ΔGbp1,2 wing discs (Fig. 6E, F). Given that cell lysis is known to release multiple active proteases, our data suggest a model in which wound detection in the *Drosophila* notum depends on latent pro-Gbps in the extracellular space, acting as protease detectors, reporting the presence of wound-induced cell lysis via Mthl10 signaling.

### Enzymatic generation of a diffusible signal explains wound-size dependence

This protease/Gbp/Mthl10 model is substantially more complex than the delayed-diffusion model of an unknown signal presented previously in Shannon et al^8^. That initial model was based on diffusive spread of an unknown wound-induced signal. Although it lacked mechanistic detail, the prior model fit the data well and provided a useful parameterization of response dynamics: a total amount of signal released compared to its detection threshold (M/Cth), a back-propagated time delay *t*_0,min_ at which calcium signals first become apparent, and an effective diffusion constant α_eff_ describing the rate at which signal spreads distally. A close look at these fitted parameters shows that the time delay and effective diffusion constant have definite trends with wound size (Fig. 7F,F’; *n* = 26 wounds with diameters > 15 μm).

**Figure 7:**
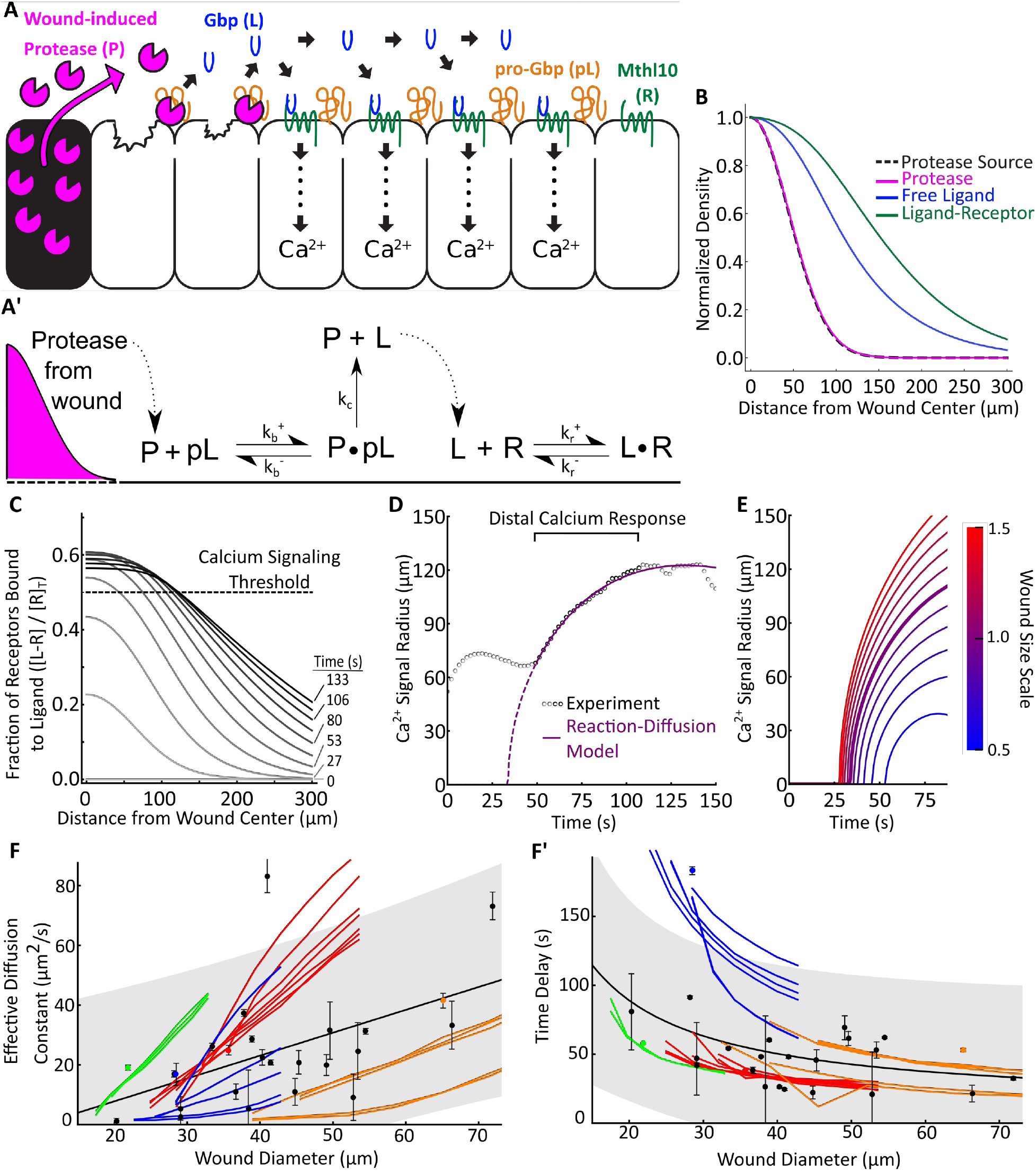
Reaction-diffusion (RD) model for epithelial wound detection. (**A, A’)** Model schematic and reactions. Diffusible proteases (P) released from lysed and damaged cells cleave immobile pro-ligand (pL), releasing diffusible, activated ligand (L) that binds to its cognate cell-surface receptor (R). **(B)** RD model output of concentration of protease (pink line), free ligand (blue line), and ligand-receptor complex (green line) as a function of distance from the wound center 1 minute after wounding. Each line is normalized by the component’s maximum concentration over all space 1 minute after wounding. The spatial extent of the protease source (black dashed line) is normalized so that its maximum is 1. (**C**) Representative model output for the fraction of receptors bound to ligand as a function of distance from the wound at various times. Threshold for triggering calcium response is taken as 0.5 (dashed line). (**D**) Calcium signal radius versus time after wounding: experimental data (open circles), and RD model fit (purple line). (**E**) Calcium signal radius from the RD model for different simulated wound sizes; each line represents the impact of scaling the wound size for an otherwise fixed set of RD parameters determined from the best fit to an individual wound response. The bold line corresponds to a scale of 1.0 and matches the best fit line in panel D. **(F-F’)** Experimental and modeled wound size dependence of the effective diffusion (F) and start time (F’) of the distal calcium response. Experimental data (black dots) are taken from n = 26 wounds larger than 15 μm in diameter. Black line and shaded region indicate the best fit curve and single prediction confidence interval respectively for a linear fit (F) and hyperbolic fit (F’) to the data. Model results (colored lines) are determined by scaling the wound size from a best fit parameter set and then parametrizing the model outputs according to a delayed-diffusion model. Each color represents a fit to the same dataset (as in Fig. S3, S4), and each line corresponds to a different best fit parameter set.

The delayed-diffusion model can describe these trends, but the description does not provide explanatory power. The trend for α_eff_ is well described as a linear increase with wound diameter *w:* α_eff_ = (0.77 ± 0.26 μm/s) *w* - (8 ± 11 μm^2^/s), whereas the trend for *t*_0,min_ is better described by a linear dependence on 1/*w*: *t*_0,min_ = (1500 ± 700 μm s) *w^-1^* + (12 ± 19 s). The coefficients of the *w* or 1/*w* term are non-zero with p-values of 0.006 and 0.04, respectively). Although one can postulate explanatory hypotheses, the delayed-diffusion model itself provides no reason for the wound-size dependence of the response dynamics. We thus wanted to explore whether the observed trends would fall out naturally from a more detailed model based on the protease/Gbp mechanism.

To do so, we constructed a computational reaction-diffusion model as outlined in Fig. 7A, A’. In brief, wound-induced cell lysis releases proteases that enzymatically cleave extracellular pro-Gbps into their active forms, which then reversibly bind Mthl10 receptors. This abstraction treats all classes of proteases as a single temporally and spatially varying protease activity and groups all Gbps into single pro- and active Gbp pools. Pro-Gbps and membrane-bound Mthl10 are treated as stationary, but released proteases and active Gbps are allowed to diffuse. The model is solved for six species – protease, pro-Gbp, protease::pro-Gbp complex, free active Gbp, receptor-bound Gbp, and free receptor (Fig. 7B) – over time and a 2D radially symmetric space. Signaling downstream of the receptor is approximated as a threshold event that releases calcium from intracellular stores when a given fraction of receptors are bound by ligand (Fig. 7C, dashed line). For complete mathematical details, see Supporting Text.

As constructed, this model has nine free parameters including protease release characteristics, initial component concentrations, diffusion constants, and reaction rates. Given that experimental response dynamics are adequately fit by a simpler model using just three parameters, one should not expect fits of the more detailed model to place strong constraints on all parameters. Instead, the detailed model falls into the category of “soft” or “sloppy” behavior common in systems biology^50^: parameters are weakly constrained, often varying over orders of magnitude, but predictions of model output are nonetheless robust and useful. In an attempt to provide stronger parameter constraints, we did try simultaneous fitting of multiple experimental data sets using a set of nine shared parameter values. Such fits did not describe the data well (Fig. S3A). As a second attempt and recognizing that wound size varied among experiments, we also tried simultaneous, multiple-data-set fits using seven shared biochemical parameter values and two experiment-specific wound parameters; however, this additional flexibility was still insufficient (Fig. S3B). We thus proceeded with fits of the full nine-parameter model to individual data sets and used the resulting parameter sets as the basis for comparing further model predictions with experimental data.

We selected four typical experimental responses and fit each calcium signal radius versus time to the reaction-diffusion model using a constrained least-squares approach (see Supporting Text). For each experimental response, we conducted 32 fits with different randomly selected sets of initial parameter guesses. We used the single best fit to estimate the variance and kept all fits for which the chi-squared statistic indicated an equivalently good fit at the 95% confidence level (3-7 fits for each experiment). A single experimental response and model fit are shown in Fig. 7D, and a full grid of all good fits is shown in Fig. S4A. As expected for a “soft” model, the superset of parameter estimates from all good fits yields distributions for individual parameters that vary by orders of magnitude (Fig. S4B).

Despite these variations among best-fit parameters, the model makes robust predictions for its output’s dependence on wound size. For each set of best fit parameters (a set being a group of nine parameters that yield a good fit), we solved the model with all parameters fixed, save two that scale with wound size: the 1/e^2^ radius of the protease source, which scales linearly with wound radius; and the total amount of protease activity released, which scales as wound radius squared. With these two parameters scaled in this manner, the peak density of the protease source (activity released per unit area) is held constant. As the example in Fig. 7E shows, increasing wound size in the model yields smaller time delays and more rapid diffusion-like signal spread. To better compare to experimental trends, we parameterized the detailed model output in the same way as experimental data, i.e., by fitting its calcium signal radius versus time to the three-parameter delayed-diffusion model. The resulting wound-size dependencies fall out as two natural predictions of the detailed model. First, the parameterization shows that the effective diffusion constant in the detailed model output increases with wound size around every one of the widely varying best-fit parameter sets (each colored line in Fig. 7F). Although individual curves flatten out as a approaches zero, the trends are mostly linear and the slopes are comparable or greater than the trend observed across all experiments. Second, the parameterization shows that the signaling time delay in the detailed model varies in a roughly hyperbolic manner with wound size. These predicted trends are also similar to experiments (Fig. 7F’).

Within the detailed model, the wound-size-dependence for the time delay and spread rate of the distal response can be traced to a generalized mechanism with two key structural features. First, the signal (i.e., active Gbp) builds up gradually over time. Second, the signal diffuses rapidly enough during this build up to spread well beyond the spatial extent of its source. The first feature by itself provides both a time delay associated with reaching threshold and a diffusion-like spread associated with the threshold boundary moving further from the wound as signal builds; however, it does not yield a wound-size dependence. Even though smaller wounds produce smaller amounts of protease activity and thus smaller integrated signals, they do so over smaller areas to yield the same signal density as larger wounds. The second key structural feature is needed to make sure that smaller or larger integrated signals are spread over comparable areas, decreasing the signal density for smaller wounds, increasing it for larger wounds, and yielding the observed wound-size dependence for α and *t*_0,min_.

### Computational model identifies key role for cell lysis over time

In the detailed model construction used here (Fig. 7A), the rate of signal spread is controlled largely by the rate of signal accumulation plus the diffusion constant of active Gbp. Diffusive spread of the protease itself is minimal (Fig. 7B). The rate of signal accumulation itself is controlled by the rate constant for the release of protease activity via cell lysis and the enzymatic rate constant for proteolytic cleavage of pro-Gbp. Among the set of best fits, these two model parameters can compensate for one another, as shown by their inverse correlation (Fig. S5). Interestingly, only a few of the best fits had a quick release of protease activity after wounding (< 1 s) and modifying the model to force an instantaneous release of this activity failed to fit the experimental data. Given that laser-induced wounds are made very rapidly – cavitation bubble generation and collapse occur within microseconds – the best-fit model’s requirement for much slower protease release was an unexpected prediction. We hypothesized that progressive cell lysis, rather than an instantaneous release of proteases from all damaged cells, may be occurring. This led us to reexamine live imaging of nuclear-mCherry-labeled cells around wounds. Although a small number of cells were destroyed immediately by the ablation process, a much larger group of surrounding cells were observed to lyse progressively over ~45 seconds (Fig. S6).

Quantitative modeling based on the protease/Gbp mechanism thus provides explanations for the distal calcium response’s wound-size dependencies and makes an unexpected prediction regarding slow protease release that matches a reexamination of experimental data. Further model predictions are provided by the sensitivity analysis shown in Fig. S7. Several parameters have a strong impact on the timing and reach of the distal calcium response, including all those related to the release of protease activity, plus the rate constant for pro-Gbp cleavage, and the initial amount of pro-Gbp present in the extracellular space. It is especially interesting that the amounts of pro-Gbps are predicted to have a strong impact on the calcium response, as these are known to vary with environmental conditions^48^. The inclusion of an enzymatic step in the wound-detection pathway provides both signal amplification and multiple options for regulation. Quantitative modeling provides a set of potentially experimentally testable predictions for how this regulation could function *in vivo.*

### Conclusion

Based on our experimental and modeling studies, we propose a mechanism by which extracellular pro-Gbps act as protease detectors that can be cleaved by multiple types of proteases released from lysed cells. Such lysis is a feature of all types of wounds, whether induced by trauma or pathogens, *e.g.,* viral lysis. Both the proteases and the newly cleaved Gbp diffuse extracellularly from the wound site allowing Gbp to bind to the GPCR Mthl10, alerting a cell within seconds to the presence of a nearby wound. Although Gbp and Mthl10 do not have direct orthologs in chordates, the cytokine and GPCR families are widely conserved. Additionally, we note that similarities exist between this mechanism and wound-defense signaling in the plant *Arabidopsis,* where damage-induced cytosolic calcium activates a cysteine protease which releases an immunomodulatory peptide^16^. Because the basic circuitry is similar across kingdoms, our study suggests an ancient strategy for wound detection based on cell lysis. Furthermore, damage or pathogen induced activation of proteins by proteolytic cleavage has been well documented in the cases of spätzle in the Toll pathway^51–53^, thrombin and fibrin in the blood coagulation pathway^54–56^, and IL-1β and IL-18 in the pyroptosis pathway^57–59^. As these examples make clear, proteases are already known to play critical roles in blood clotting and immune signaling, and this study finds that proteases are instructive signals in epithelial wound detection.

## Acknowledgements

We thank E. Baehrecke (University of Massachusetts Medical School, Worcester, MA) for providing the *UAS-IP_3_* sponge fly stock; T. Koyama (IGC, Lisbon, Portugal) for providing the *ΔGbp1,2* fly stock; the Bloomington *Drosophila* Stock Center (Bloomington, IN) and the National Institute of Genetics (Shizuoka, Japan) for *Drosophila* stocks; G. Struhl (Columbia University, New York, NY) for providing the plasmid containing the *Actin5c* promoter; G. Neuert, M. Tyska, J. Nordman, and I. Macara (Vanderbilt University, Nashville TN) for comments on the manuscript. Funding: This work was supported by the National Institute of General Medical Sciences (1R01GM130130 to A.P.M. and M.S.H.), the National Institute of Arthritis and Musculoskeletal and Skin Diseases (R21AR072510 to A.P.M.). E.K.S. was supported by the National Institute of Cancer (5T32CA119925) and J.O.C. was supported by the National Institute of Child Health and Human Development (T32HD007502) and the American Heart Association (19PRE34410069 to J.O.C.). Author Contributions: All authors contributed to overall project design. J.O.C., E.K.S., K.S.L., M.S.H., and A.P.M. designed *Drosophila* experiments. K.S.L. and N.N. created the null mutant lines. J.O.C. and E.K.S. performed pupal wounding experiments. J.O.C. and F.B.A. performed larval wing disc experiments. A.C.S. and M.S.H. designed and created the computational model. J.O.C., A.C.S., M.S.H., and A.P.M. wrote the manuscript.

## Supplement to

**Figure S1:**
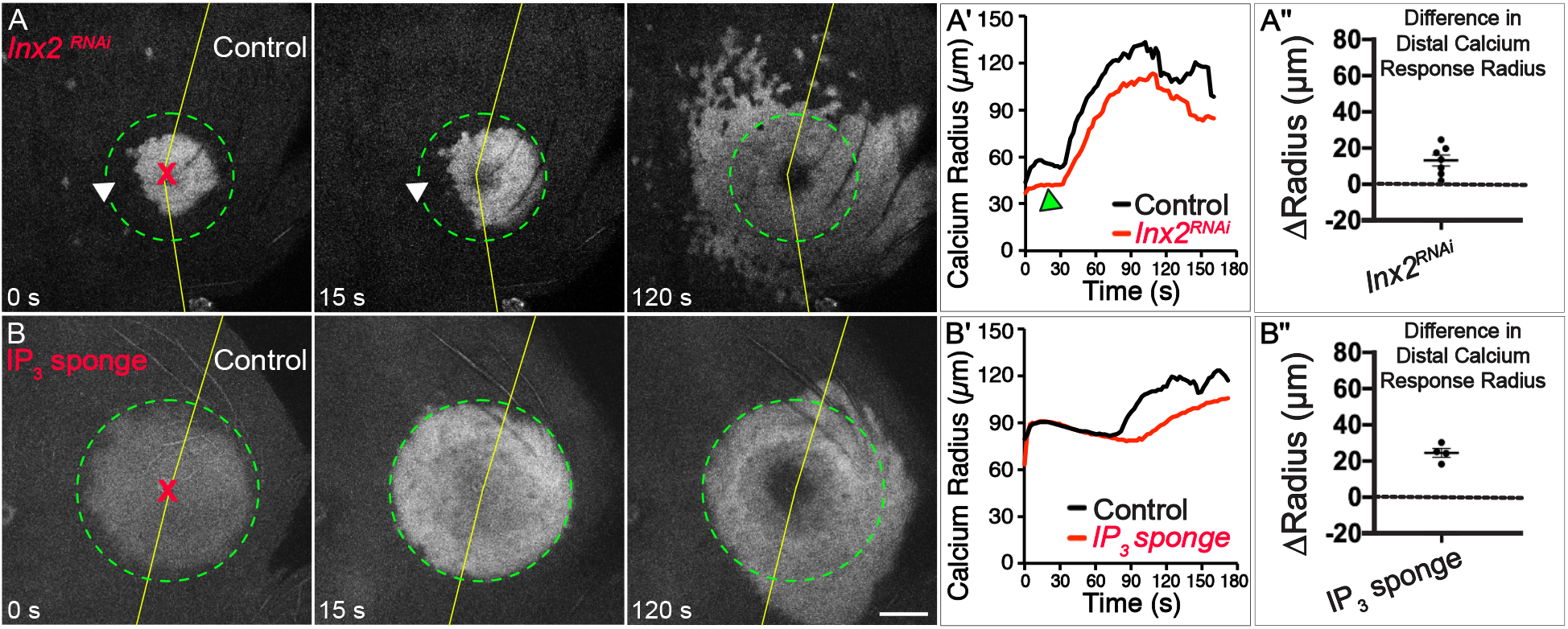
Wounds signal calcium to proximal cells through gap junctions and to distal cells through IP_3_. (**A-B**) GCaMP6m reporter showing cytosolic calcium in representative samples of various knockdowns in the *Drosophila* notum. Gene manipulations are performed in the *pnr* expression domain (left side) and wounds (red X) are administered at the *pnr* domain boundary (yellow line). Maximum extent of first expansion is marked by dashed green circle. Scale bar = 50 μm. (**A**) Knockdown of gap junctions (*Inx2*^RNAi^) attenuates first calcium expansion (arrowheads); the distal calcium response still occurs in the knockdown domain, but with a “speckled” pattern, as reported previously^8,32,36^. Although the *Inx2* knockdown appears to cause a slight reduction in calcium radius (A’–A’’), the speckled nature of the signal would cause the automated algorithm that identifies the radius to underestimate it. (**B**) Expression of an IP3 sponge has no effect on the first calcium expansion, but the distal calcium response is delayed. (**A’’–B’’**) Difference in calcium signal radii (ΔRadius) at maximum extent of the distal calcium response for control minus pnr sides of each wound; pnr-side genotypes indicated.

**Figure S2:**
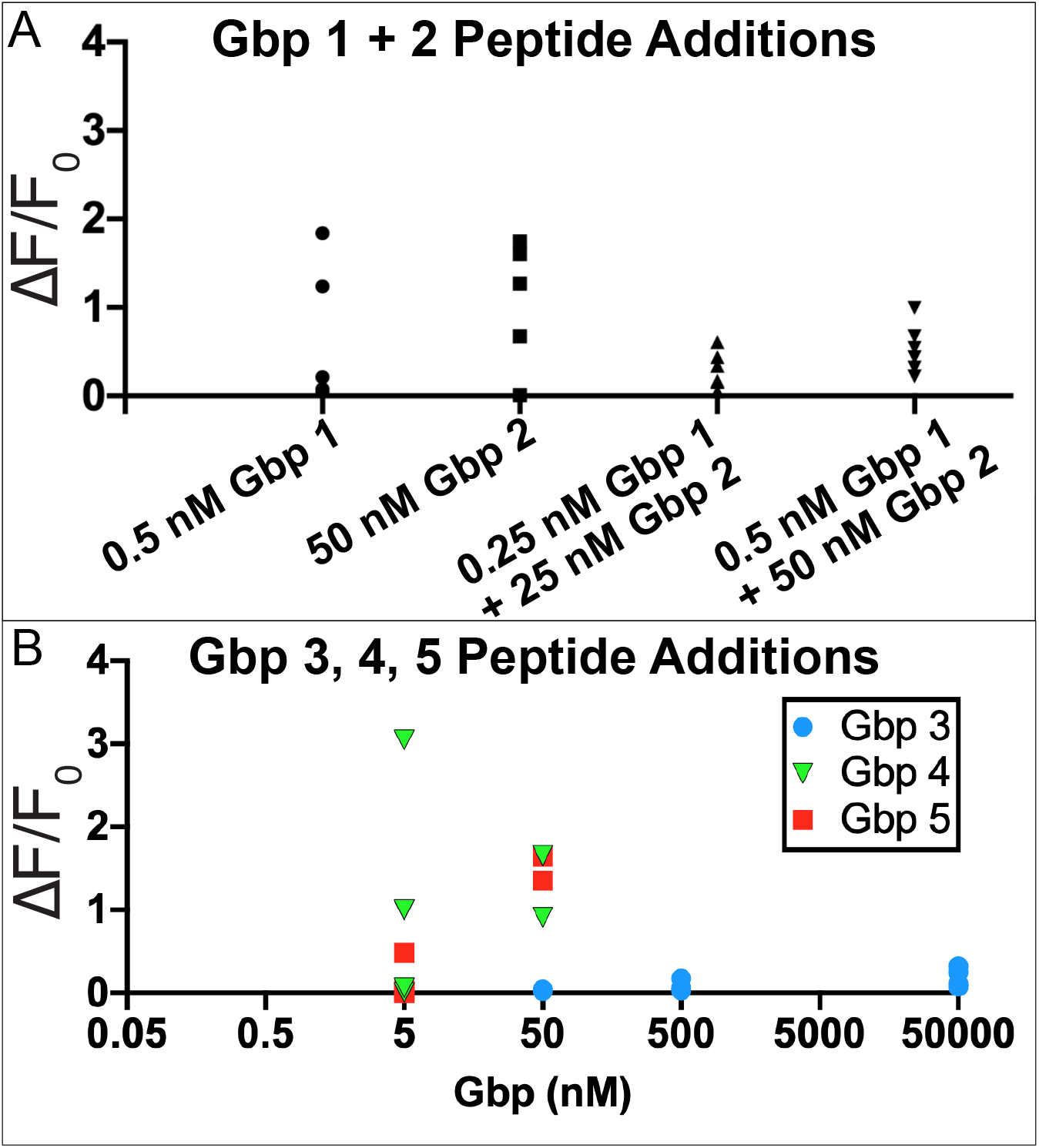
Gbps 1 and 2 do not act synergistically, and Gbps 4 and 5 are also sufficient to elicit calcium responses in wing discs. (**A**) Change in GCaMP6m calcium fluorescence when the given concentration of each peptide is added to *Drosophila* wing imaginal discs. The lowest concentrations of Gbp1 and Gbp2 sufficient to elicit calcium from Fig. 4 (0.5 nM and 50 nM, respectively) are shown. The calcium response is not increased when half this concentration of each (0.25 nM Gbp1 + 25 nM Gbp2) nor all (0.5 nM Gbp1 + 50 nM Gbp2) are added simultaneously, showing Gbp1 and Gbp2 do not act synergistically. (**B**) Change in GCaMP6m calcium fluorescence when a given concentration of Gbp −3, −4, or −5 peptides is added to *Drosophila* wing imaginal discs. Even at 50 μM, Gbp3 (blue circle) does not elicit a calcium response. Gbp4 (green triangle) elicits an inconsistent response at 5 nM, and consistently when added at 50 nM. Gbp5 peptide elicit a calcium response at 50 nM.

**Figure S3:**
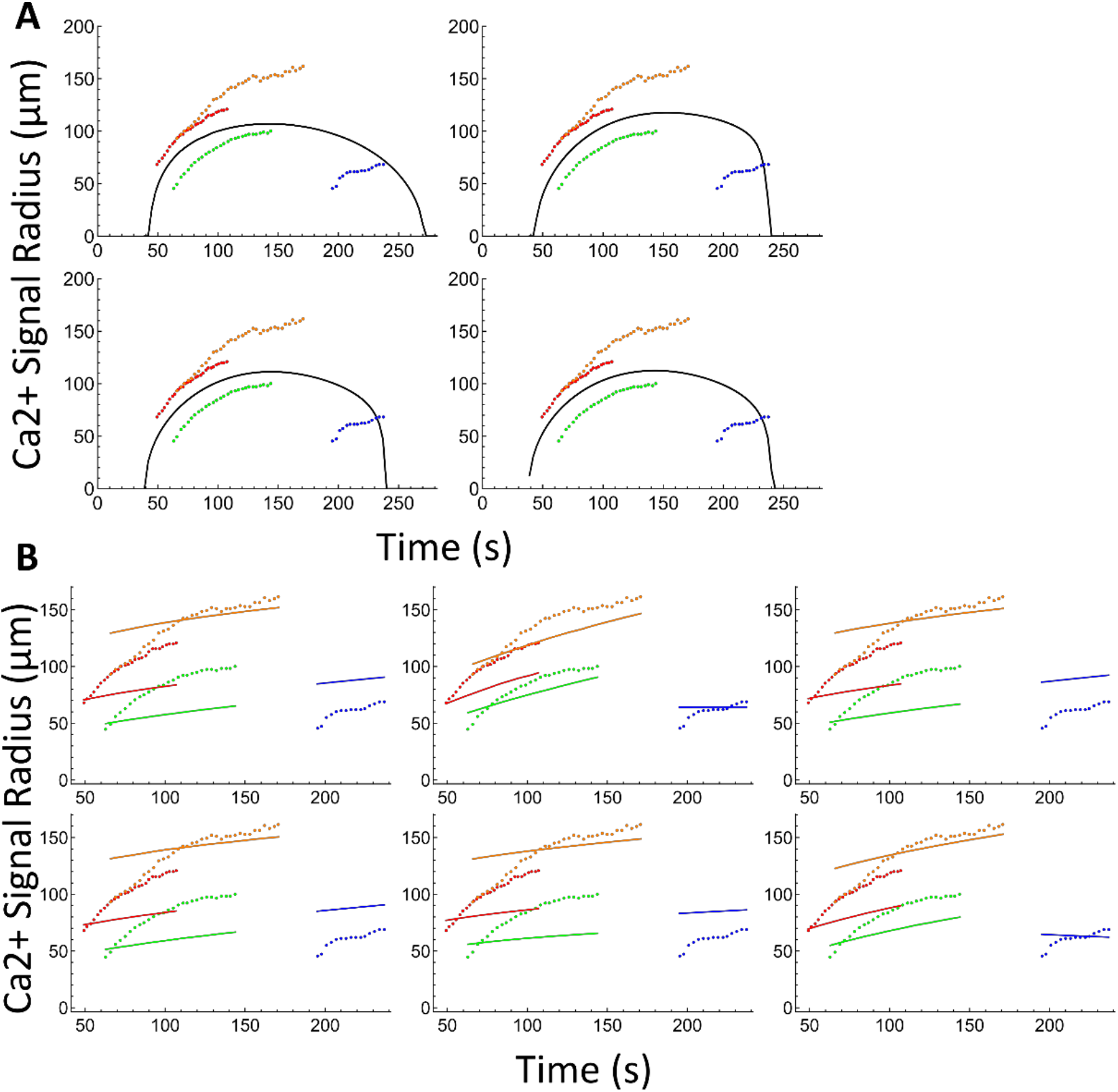
Results of simultaneously fitting the reaction-diffusion model to four selected experimental samples. (**A**) Four results of running the fitting process trying to minimize the cost function for all data sets simultaneously with a single parameter set. Model output is given by the black line, and distal calcium response data (colored points) are color coded based on the colors in Fig. S4. (**B**) Six results of running the fitting process trying to minimize the cost function for all data sets simultaneously with a single base parameter set that is scaled according to the wound size of each distal calcium response (See “Distal calcium response properties from the model” supplement section for specifics on scaling parameters based on wound size). Model output (lines) and distal calcium response data (points) are color coded based on the colors in Fig. S4. Colored lines correspond to model output given the wound size of the same colored distal calcium response data points.

**Figure S4:**
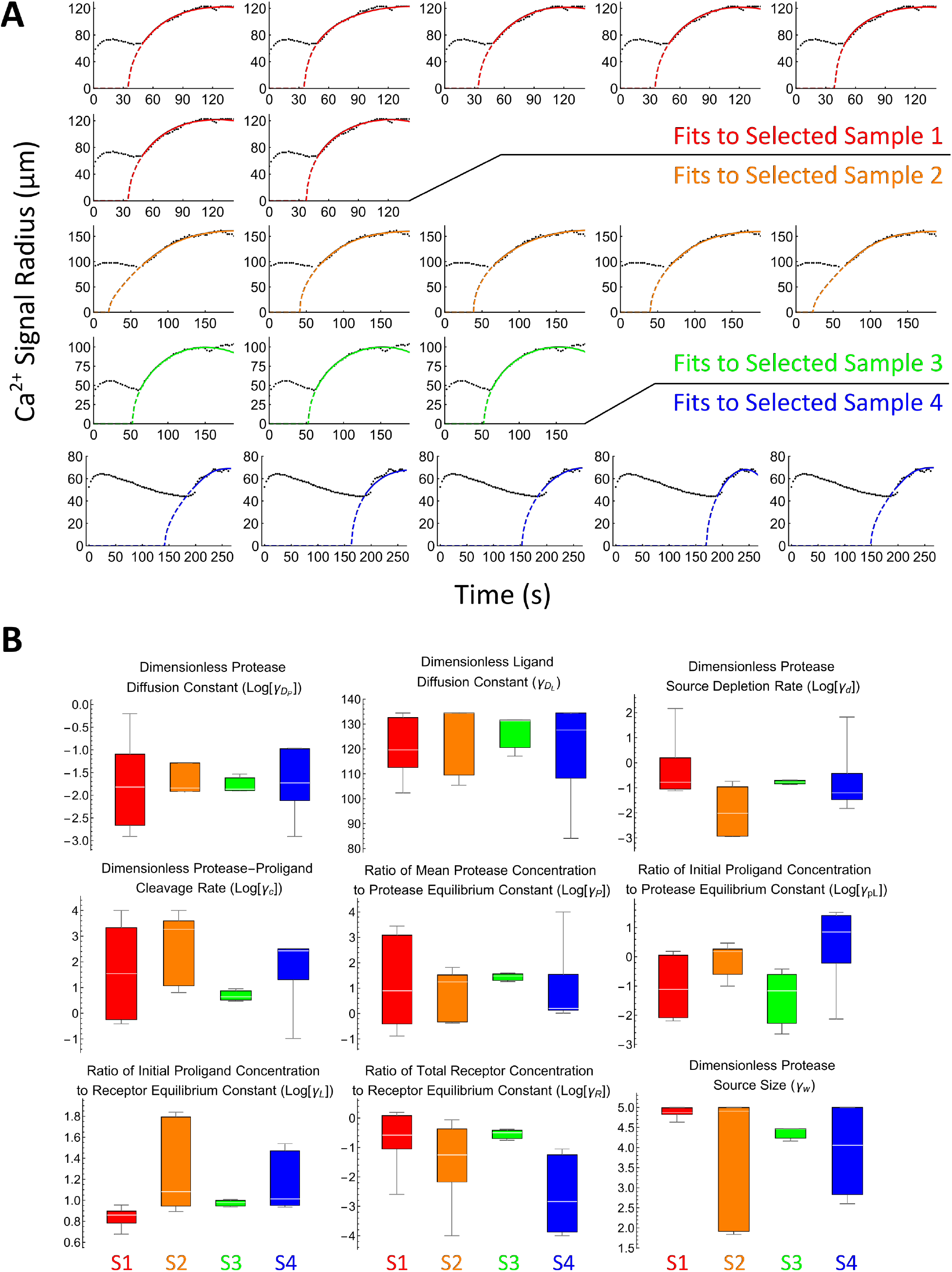
Results of fitting the reaction-diffusion model to four selected experimental samples. (**A**) Calcium signal radius versus time after wounding: experimental data (black circles), and RD model fits (colored lines). Each graph is a different best RD model fit to the distal calcium response data. (**B**) Box and whisker plots of best fit parameters to each selected distal calcium response. Parameters are defined further in the Supplemental Text. Colors correspond to the colors in A. Note that some of the plots correspond to the logarithm of the parameters to emphasize spreads across orders of magnitude.

**Figure S5:**
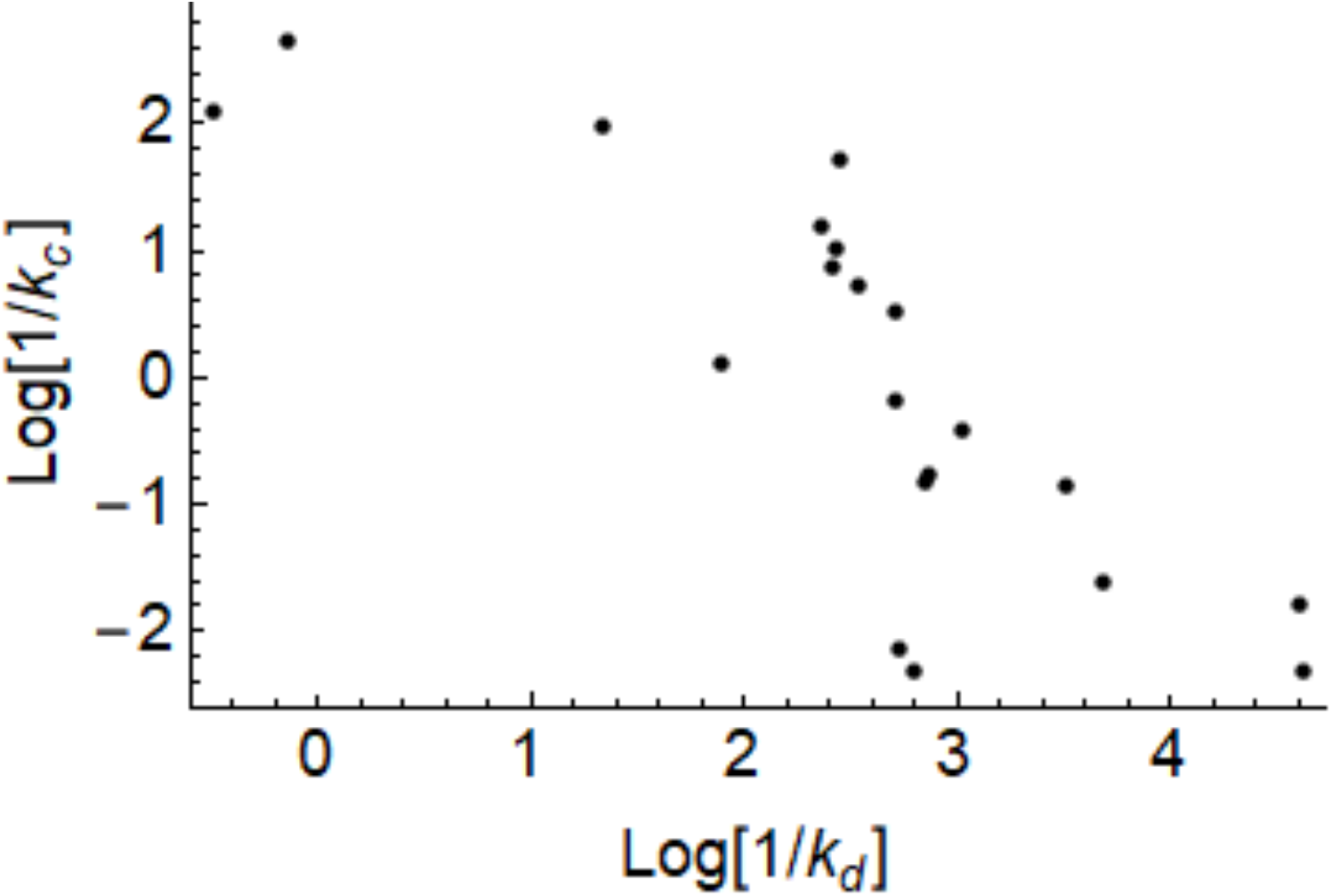
Correlation of time constant parameters. Time constant of pro-ligand cleavage (1/k_c_) versus time constant of protease release and activation (1/k_d_).

**Figure S6:**
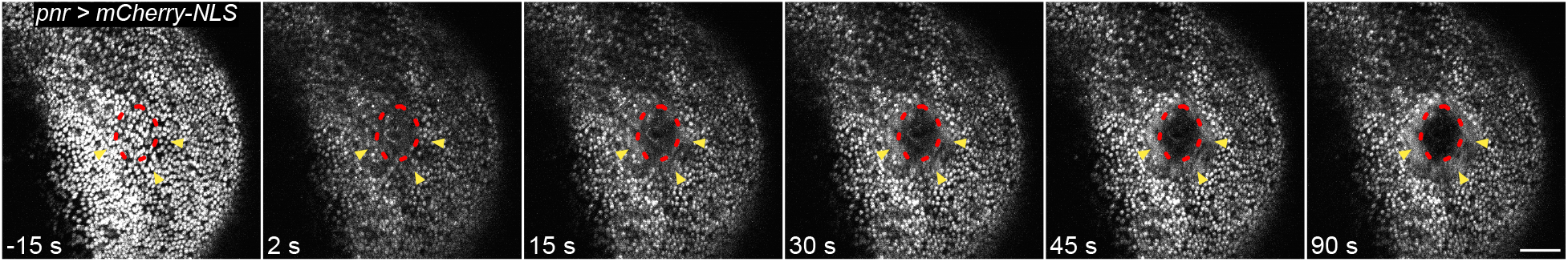
Cell lysis occurs over time after a single laser wound. Over the first 90 s following wounding, severely damaged cells at the center of the wound completely lose fluorescence (red outline), indicating cell lysis. The black region devoid of signal at 90 s corresponds to the region of fully lysed cells at the center of a wound. This region of cell lysis increases over 45-90 seconds after wounding. Yellow arrowheads indicate cells with damaged nuclear membranes, which release nuclear-localized mCherry from nuclei into the cytosol. Scale bar = 50 μm.

**Figure S7:**
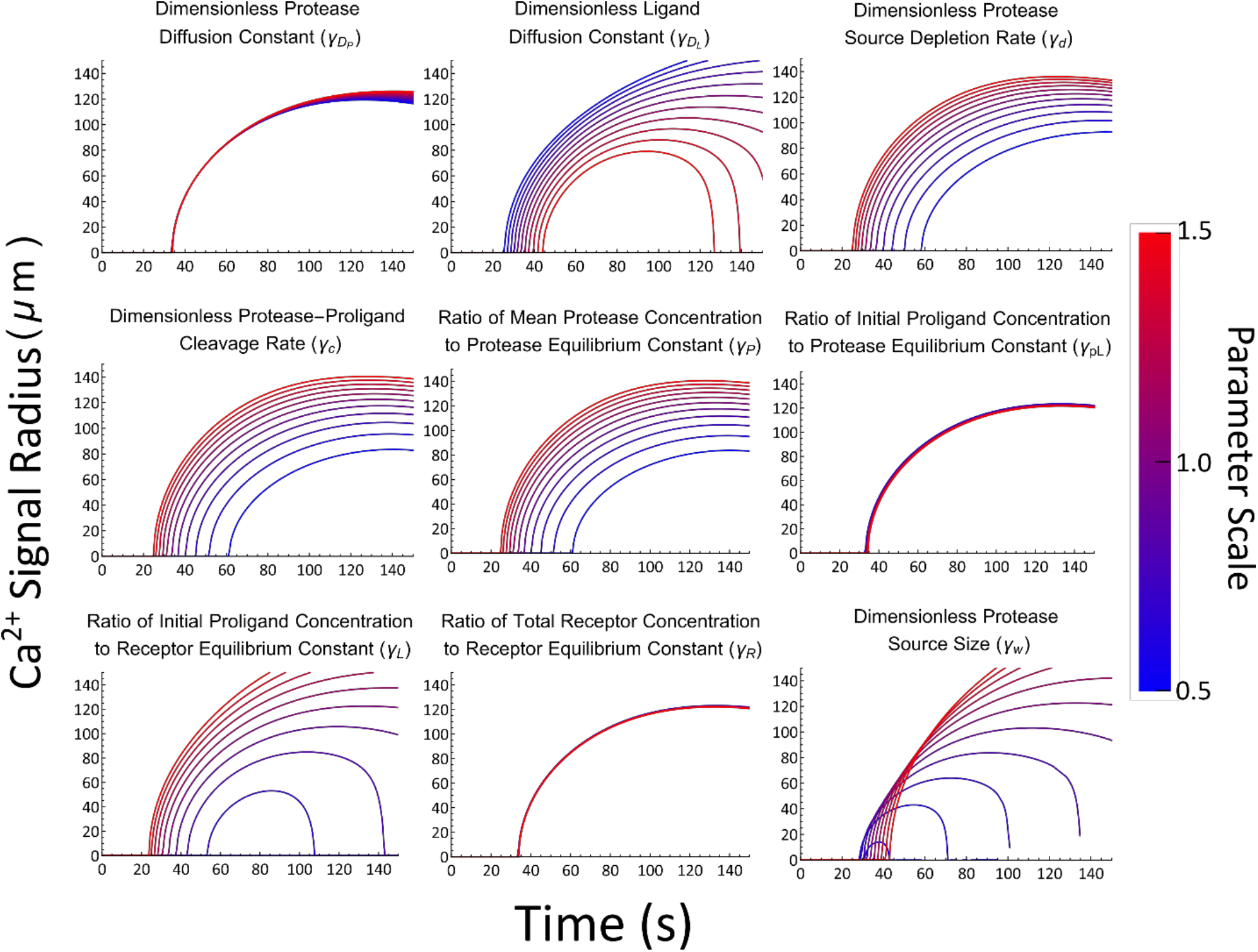
Parameter variation effect on RD model output. Calcium signal radius when varying a single parameter from a set of RD parameters determined from the best fit to an individual wound response (Purple line, Fig. 7D). Parameter scale indicates by how much the corresponding parameter was scaled to obtain lines of the corresponding color.

## Supplementary Materials and Methods

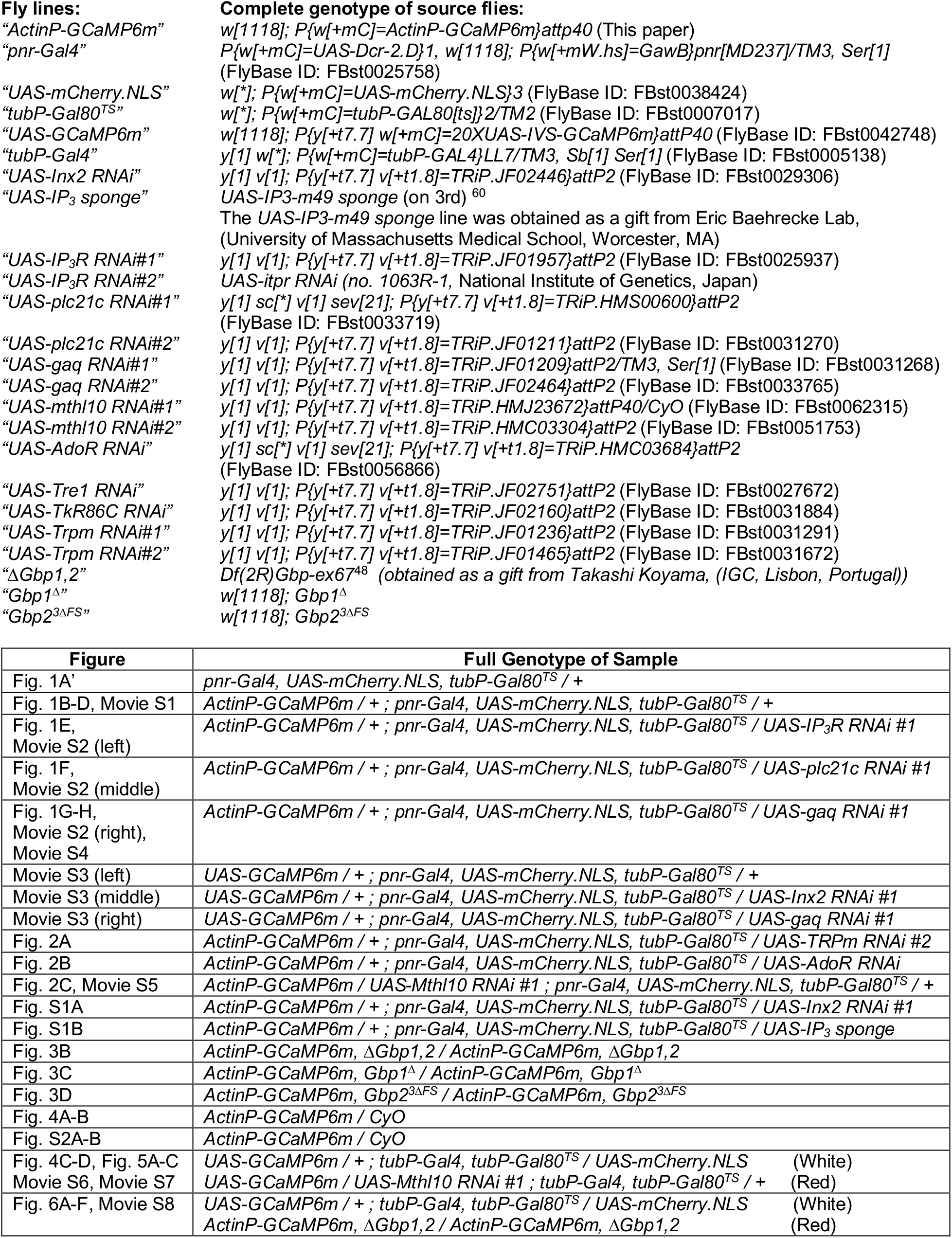

### *Drosophila* husbandry

*Drosophila* crosses were maintained at 18°C for 2 days to inhibit Gal4 activation in the presence of Gal80^ts^ during embryogenesis. Progeny were then incubated at 29°C to activate Gal4, where they remained until experimentation. Thus, 3^rd^ instar larvae were incubated at 29°C for 3-4 days before wing disc dissections and pupae were incubated at 29°C for 4-5 days before wounding.

### Pupal mounting

Pupal mounting was performed as described previously^8^. White prepupae were identified and aged for 12-18 hours After Puparium Formation (APF) at 29°C. Multiple pupae were placed on a piece of double-sided tape (Scotch brand, catalog #665), ventral side down on a microscope slide, and their anterior pupal cases were removed with fine tipped forceps to reveal the epithelium of each notum (as in Fig. 1A). The entire piece of double-sided tape (with dissected pupae) was gently lifted from the microscope slide and adhered to a 35 mm x 50 mm coverslip (Fisherbrand, cat#125485R) so the pupal nota were laid against the coverslip, with the pupae between the coverslip and the tape layer. Then, an oxygen permeable membrane (YSI, standard membrane kit, cat#1329882) was applied over the pupae and secured to the coverslip with additional double-sided tape so pupae would not become dehydrated or deprived of oxygen.

### Pupal survival

When mounted as described above, pupae developed normally for the next 3-4 days until eclosion, whereupon the adult flies crawled out of their cases and became stuck on the double-sided tape a few inches away. Pupal survival was measured as whether a pupa had developed into an adult fly and emerged from its case in this manner within 7 days after wounding (to ensure a pupa delayed in development was not accidentally miscounted as dead). Slides of pupae were returned to 29°C during this recovery phase after wounding to maintain Gal4 activation.

### Live imaging

Live imaging of pupae was performed using a Zeiss LSM410 raster-scanning inverted confocal microscope with a 40X 1.3 NA oil-immersion objective. Raster-scans were performed with a 2.26 seconds scan time per image with no interval between scans. Live imaging of wing discs was performed on the same scope, with the confocal settings turned off to maximize imaging depth, using a 25X 0.8 NA air objective. Raster-scans were performed with a 2.26 s scan time per image with a 15 s interval between scans. Notum picture in Fig. 1A’ was taken on the same scope at 10X 0.50 NA air objective.

### Laser ablation

Laser ablation was performed using single pulses of the 3rd harmonic (355 nm) of a Q-switched Nd:YAG laser (5 ns pulse-width, Continuum Minilite II, Santa Clara, CA). Laser pulse energies were on the order of 1 μJ, but were varied day to day and based on focal plane of ablation in order to optimize consistent wound sizes. A separate computer-controlled mirror and custom ImageJ plug-in were used to aim and operate the ablation laser so that ablation could be performed without any interruption to live imaging. The frame during ablation was retroactively considered t = 0 s.

### Puncture wounding

Multiple pupae were placed on a piece of double-sided tape (Scotch brand, catalog #665), ventral side down on a microscope slide, and each anterior pupal case was removed with fine tipped forceps to reveal the notum epithelium (as in Fig. 1A). The slide was mounted on the stage of an upright epifluorescence microscope (Zeiss Axio M2) and imaged with a 1 s interval on a 5x objective (Zeiss EC Plan-NEOFLUAR 420330-9901). Pupae expressing GCaMP6m were manually punctured with an electrolytically sharpened tungsten needle (Fine Science Tools, No:10130) while imaging. The frame when the puncture occurred was retroactively considered t = 0 s. Unfortunately, many puncture wounds resulted in a bubble of hemolymph that pooled over the wound, obscuring the initial influx and first expansion from multiple samples. However, the distal calcium response was always speckled in gap-junction knockdowns and always absent in Gq-pathway knockdowns, recapitulating laser wounds.

### Wing disc mounting

Wing discs from 3^rd^ instar larvae were dissected in Schneider’s *Drosophila* media (Gibco, Life Technologies, Ref:21720-024) and immediately mounted in a small bubble of Schneider’s *Drosophila* media on coverslips (Fisherbrand, cat#125485R) for imaging. A pap pen (RPI, catalog #50-550-221) was used to trace a hydrophobic barrier around the wing discs on the coverslip. Two wing discs, control and experimental, were mounted side-by-side in the same media bubble. The control disc was identifiable by the presence of mCherry which was not present in the experimental disc. Images were taken to establish a baseline of GCaMP fluorescence, and then potential calcium activators were pipetted directly into the media bubble over the wing discs. The concentration of calcium activators in the text refers to the final concentration after addition to the media bubble. The image taken during pipetting was retroactively considered t = 0 s.

### Drosophila extract preparation

Fly and larval extract was made in a similar manner to that described previously in ^61^ and originally in ^62^. For adult fly extract, 100 healthy adult flies with a female:male ratio of 3:1 were homogenized in 750 μl of Schneider’s *Drosophila* media. Similarly, for larval extract, 100 healthy 3rd instar larvae, from non-overcrowded bottles, were homogenized in 750 μl of Schneider’s *Drosophila* media. The resulting homogenates were centrifuged at 4°C for 20 minutes at 2600*g*. The supernatant was heat treated at 60°C for 5 minutes, before a final centrifugation at 4°C for 90 minutes at 2600*g*. The final supernatant – considered 100% extract – was aliquoted and stored at −20°C.

### Protease Inhibition Experiments

To test whether *ΔGbp1,2* Larval Extract elicits a calcium response in control wing discs via proteases, we pre-mixed protease inhibitor cocktails or vehicle-only controls either into the extract or media bubble before addition to the wing disc. Specifically, we used 1) Cell Signaling Technologies Protease inhibitor cocktail (Cell Signaling Technology, #5871S), which had no effect on the extract-mediated calcium response of *ΔGbp1,2* Larval Extract when mixed at or below the 2x recommended concentration; at 3x recommended concentrations, the inhibitor itself elicited an ectopic calcium response on the discs, making it unusable for properly testing the extract-mediated calcium response. 2) MS-Safe Protease and Phosphatase inhibitor (Millipore Sigma, MSSAFE), which had no effect on the extract-mediated calcium response of *ΔGbp1,2* Larval Extract when mixed at or below the 1x recommended concentration; at 1.5x recommended concentrations, the inhibitor itself elicited an ectopic calcium response on the discs, making it unusable for properly testing the extract-mediated calcium response. 3) Two other protease inhibitors tested (Millipore Sigma, 539134 and Millipore Sigma, 539133) used DMSO as a vehicle, which itself induces an ectopic calcium response in wing discs at even small concentrations (1% final v/v). Therefore, both of these protease inhibitors were not unusable for properly testing the extract-mediated calcium response.

### Peptide synthesis

The amino acid sequence for Growth-blocking peptides 1–5 is shown below, as described previously.^43^

Gbp1 (CG15917): ILLETTQKCKPGFELFGKRCRKPA

Gbp2 (CG11395): SLFNLDPKCAEGLKLMAGRCRKEA

Gbp3 (CG17244): MVAMIDFPCQPGYLPDHRGRCREIW

Gbp4 (CG12517): ILLDTSRKCRPGLELFGVRCRRRA Gbp5 (CG14069): MLLEIQKRCWAGWGLLAGRCRKLA These sequences were sent to Genscript (Piscataway, NJ. USA) for peptide synthesis under conditions to maintain the disulfide bond. The lyophilized peptide was reconstituted in ultrapure water, diluted to a concentration of 0.1 μM, aliquoted, and frozen at −80°C.

### Protease preparation

Cysteine protease Clostripain (Alfa Aesar, Thermo Fisher, Stock: J61362) was reconstituted in a TBS solution (10 mM Tris, 1 mM CaCl2, 50 mM NaCl, 2.5 mM beta-mercaptoethanol) to a final concentration of 1mg/mL (18.9 μM). This was aliquoted and frozen at −20°C. Serine protease trypsin (Gibco, Life Technologies, Ref: 25300-054) arrived at a concentration of 0.05% w/v (21.4 μM) and was refrigerated at 4°C.

### *ActinP-GCaMP6m* generation

*pBPw.Act5CP-GCaMP6m* was generated from the GCaMP6m plasmid pGP-CMV-GCaMP6m (Addgene, originally in ^63^) and the Actin5c promoter ^64^. Importantly, this construct features the full 4.4 Kb genomic enhancer sequence for actin, containing all the regulatory elements to drive ubiquitous expression, rather than the more commonly used 2.6 Kb actin promoter sequence which is not highly expressed in the pupal notum. This 4.4 Kb promoter was obtained as a gift from Gary Struhl (Columbia University, New York, NY). The *pBPw.Act5CP-GCaMP6m* construct was injected by BestGene (Chino Hills, CA. USA) into *Drosophila* using PhiC31 integrase at the attP40 site at 25C6 on Chromosome 2, generating the *ActinP-GCaMP6m* fly line.

### Calcium intensity analysis

Fluorescence intensity of wing discs was measured using the Measure Stack plug-in in ImageJ. The region of interest was defined around the entire wing disc using the polygon selection tool. The mean fluorescence intensity for each disc was graphed in Microsoft Excel as a function of time, with 0 s corresponding to the frame when calcium activators were added to the wing disc media. The ΔF/F_0_ value for each experiment was defined as the disc’s fluorescence intensity in the initial frame of the movie subtracted from the disc’s maximum intensity, normalized to the initial frame intensity.

### *Gbp1^Δ^* generation

The *Gbp1^Δ^* null mutant was created by targeting two CRISPR-mediated double-stranded breaks to the *Gbp1* gene locus (www.crisprflydesign.org). The following gRNAs were chosen:

#1) ATTTGCTCCCATCATTTATC

#2)CGGAAAACGATGCAGAAAGC gRNAs sequences with extensions to allow for BbsI digestion were synthesized as singlestranded DNA oligos and annealed to form overhangs, then ligated to a pCFD5 vector (Addgene, #73914) digested with BbsI (NEB, #R3539S). Both plasmids were sequence-verified then injected by BestGene (Chino Hills, CA. USA) into *vas-Cas9* expressing *Drosophila* embryos (Bloomington Stock 51324). *vas-Cas9* was crossed out of progeny and mutants were identified by PCR for the presence of a *Gbp1* deletion.

On sequencing, *Gbp1^Δ^* was found to be missing 282 nucleotides, spanning the coding region corresponding to 94 amino acids from *£422* to K116. This includes the C-terminal active peptide of the Gbp 1 protein, whichspansI95 to A118.

### *Gbp2^3ΔFS^* generation

The *Gbp2^3ΔFS^* null mutant was created by targeting two CRISPR-mediated doublestranded breaks in the *Gbp2* gene locus, using the protocol above. The following gRNAs were chosen:

#1) GAATATTCAACGCTGCTG TT

#2) AATTCCATACAACCGCGTCC

On sequencing, the *Gbp2^3ΔFS^* allele was found to create multiple lesions: 8 missense mutations within the protein coding region generating the following amino acid substitutions: P41S, N42H, V51G, I61N, V94G, P150S, N192I, and S218P. Additionally, 3 small regions were deleted from the protein coding region: 30 nucleotides corresponding to Q236–Q246, 60 nucleotides corresponding to S252-T272, and 128 nucleotides corresponding to G284–R326. The last deletion induced a frameshift, creating multiple missense mutations and 4 stop codons in the final 391 nucleotides of the gene, thus preventing transcription of the region corresponding to the C-terminal active peptide of the Gbp2 protein, which spans S433 to A456.

### Calcium signal radius analysis

Calcium radius was analyzed as described previously^8^ and illustrated in Fig. S8. Briefly, to quantify the spread of calcium signals from full-frame time-lapse images, the ImageJ Radial Profile Angle Plot plug-in was used on each image to determine the average GCaMP6m intensity profile as a function of distance from the center of the wound. A custom MATLAB script was then used to determine the distance from the wound at which the intensity dropped to half its maximum (Fig. S8B,B’). This distance corresponds to the radius of the radially-averaged calcium wave and plotting radius for each video frame yields a graph of calcium signal expansion over time, which was graphed in Microsoft Excel (Fig. S8A’).

**Fig. S8.**
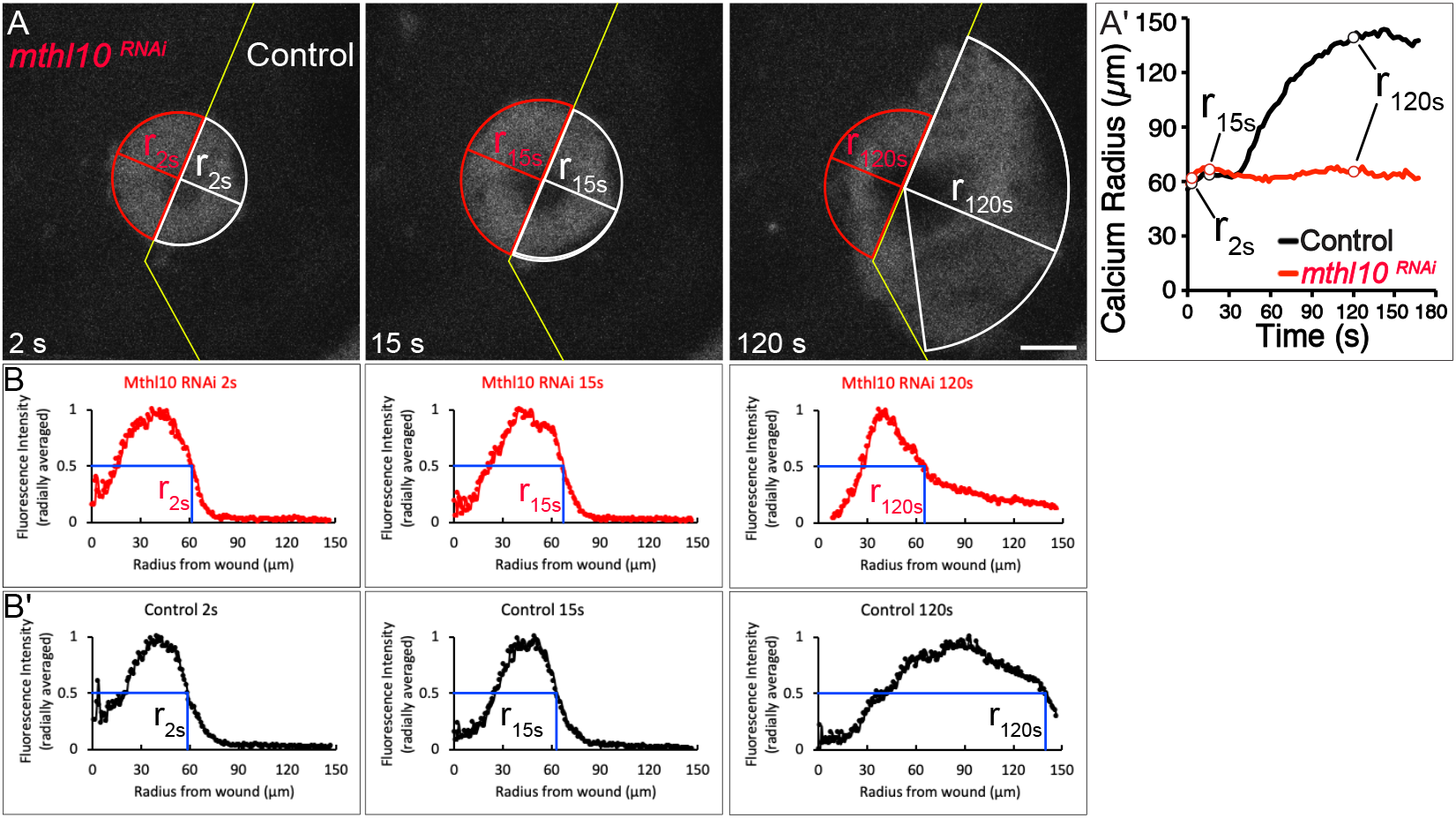
Radial analysis of calcium signaling.

For all movies with an internal control, mCherry expression was used to define the experimental and control domains. The average calcium signal within each domain was measured separately. After plotting calcium radius with respect to time, the ΔRadius value for each sample was defined as the maximum value of the control domain calcium radius minus the calcium radius of the *pnr-*expressing experimental domain at the same time point (as shown in Fig. 1E’).

### Statistical analysis

All statistical analysis was performed in Graphpad Prism. ΔRadius values for all samples of each genotype were graphed in Prism, and statistical analysis was performed by one-way ANOVA with multiple comparisons of each genotype with respect to the control group. Similarly, ΔF/F_0_ values for all wing disc experiments were graphed in Prism, and statistical analysis was performed by Student’s t-test in all cases except for Fig. 6C which was analyzed by two-way ANOVA with multiple comparisons of each genotype with respect to the control extract + control disc group. Each scatterplot displays the value for each sample as a point, with bars representing mean and SEM.

## Reaction-Diffusion Model

### The extracellular space

While the extracellular space in the pupal laser wound experiments is a threedimensional space, arguments can be made to reduce it to a one-dimensional space. First, a cuticle that exists just above the tissue essentially limits the extracellular space to a twodimensional plane. Second, due to the radial symmetry of the distal calcium response around the wound, we can assume that the system is radially symmetric about the wound center. Therefore, the relevant molecular components in the model can be assumed to only depend on two dependent variables: one spatial variable *r* that is the distance from the center of the wound, and one temporal variable *t* that is the time since wounding.

### Protease Source

Since the extent of the damage decreases farther from the wound center, it is reasonable to assume that the source of active protease follows this same trend. Therefore, the active protease source initiated by the wound is taken to be a Gaussian in space whose maximum is at the wound center. The source is also taken to decay exponentially in time to represent the depletion of protease from the total amount released from the wound. The protease source term then takes the form

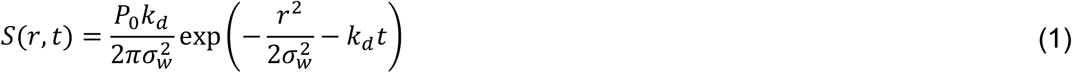

where *P*_0_ is the total number of protease molecules released from the source, *k_d_* is the rate constant for the source depletion, and *σ_w_* determines the spatial extent of the source.

It is worth noting that, while active protease could arise from various mechanisms, the model remains agnostic as to how we get to a final, activated form of the protease. Therefore, the Gaussian form of the protease source just represents where we would expect to find active protease after various activation mechanisms have occurred.

### Protease-proligand model

The protease-proligand interactions are modeled as a simple reaction, where one protease molecule reversibly binds to one proligand molecule to form a complex. This complex is then irreversibly converted into a protease molecule and a ligand molecule upon cleavage of the proligand. These processes can be modeled with the scheme shown in Fig 7A’ with rate constants 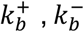 and *k_c_*.

### Ligand-Receptor model

The ligand-receptor interactions are modeled as a simple reaction, where one ligand molecule reversibly binds to one receptor molecule to form a complex. This process can be modeled with the scheme shown in Fig 7A’ with rate constants 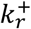 and 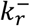. While the ligandreceptor dynamics upstream of calcium signaling can be modeled in a more detailed way, as in ^65^, these processes most likely do not influence the calcium signals on the timescale of the distal calcium response, so they are not considered here.

## Reaction-Diffusion Model

### Set of Equations

Using the law of mass action, the model reactions above can be turned into a set of coupled partial differential equations. The model assumes that all molecular components exist in a radially symmetric, two-dimensional plane where protease and ligand can diffuse with diffusion constants *D_P_* and *D_L_* respectively. All other molecular components are assumed to be immobile. This leads to the following set of six differential equations

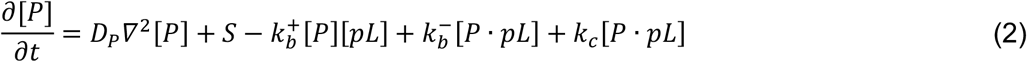

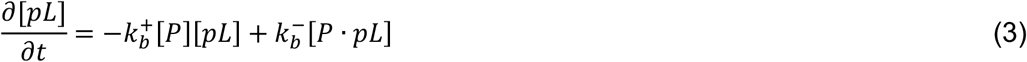

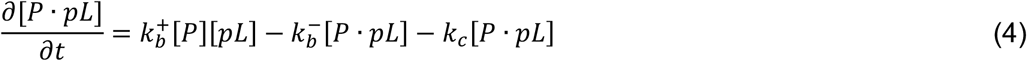

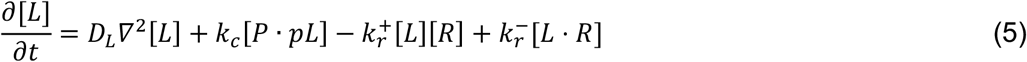

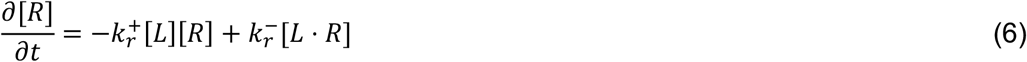

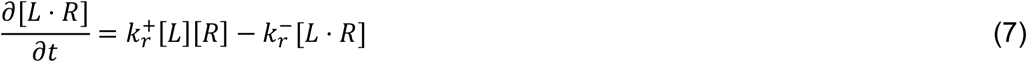

where brackets denote the concentration of a molecular component in terms of number of molecules per unit area, and the Laplacian operator ∇^2^ is expressed in terms of the polar coordinate *r* as 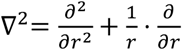.

In order to simplify this set of equations, it is assumed that ligand production is described by Michaelis-Menten kinetics and that ligand-receptor binding is at equilibrium on fast timescales. Therefore, we have

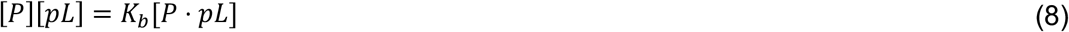

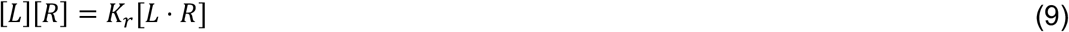

where 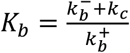 and 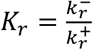. Additionally, by defining the total proligand concentration as [*pL*]_*T*_ = [*pL*] + [*P* · *pL*] and the constant total receptor concentration as [*R*]_*T*_ = [*R*] + [*L* · *R*], we obtain the following equations:

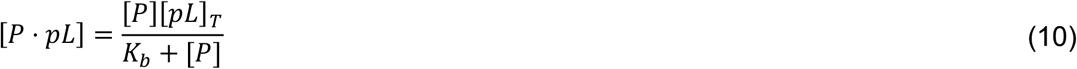

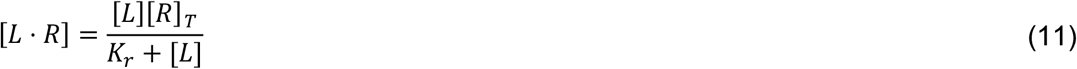

By adding together specific pairs of equations from (2) – (7) we obtain

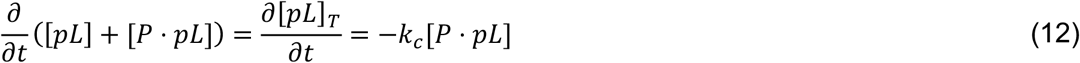

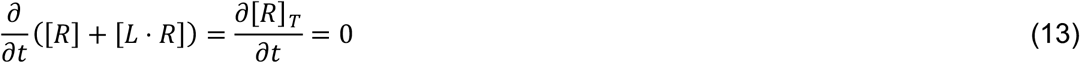

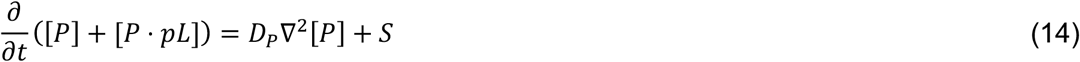

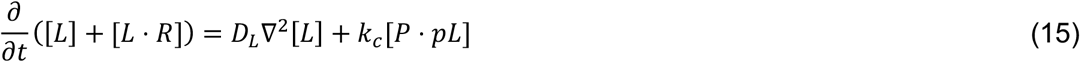

Then, by combining equations (10) – (15)we end up with a set of three differential equations and four algebraic equations

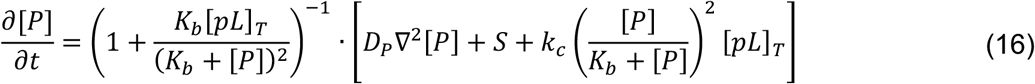

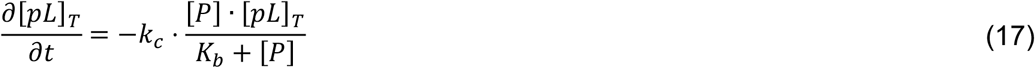

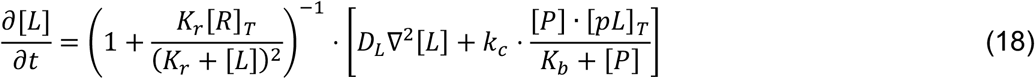

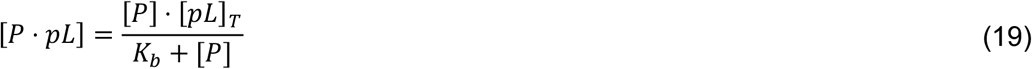

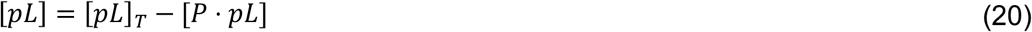

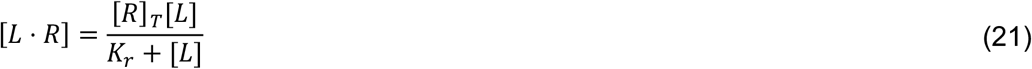

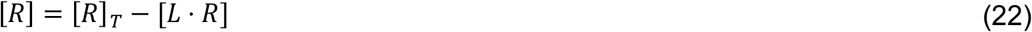

### Initial and Boundary Conditions

Due to diffusive terms in the equations, the mobile molecular components require both boundary conditions and initial conditions, whereas the immobile molecular components only require initial conditions. We assume that there is initially no protease or ligand in the extracellular space, and that a uniform proligand concentration [*pL*]_0_ is present in the extracellular space. This gives the following initial conditions

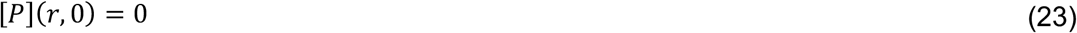

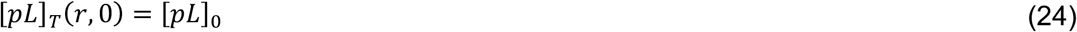

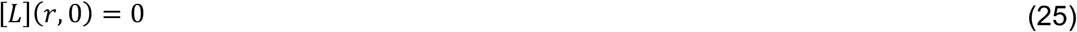

The initial conditions of the other molecular components can be obtained from the algebraic equations above, although these are not important in solving the differential equations.

For each mobile molecular component, the same boundary conditions are applied: far from the wound the concentrations should go to 0, and in order to impose radial symmetry there needs to be no flux at the origin. This leads to the following boundary conditions

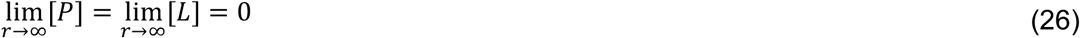

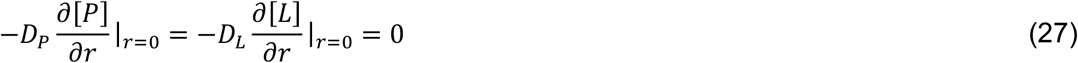

### Non-dimensionalization

To reduce the number of parameters in the model, as well as simplify the model further, the independent variables are scaled in the following way:

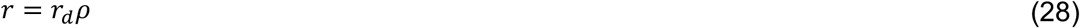

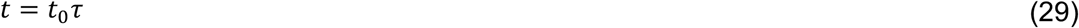

where *r_d_* = 10 *μm* is the average ablation radius following wounding, *t*_0_ = 47 *s* is the median time after wounding before the distal calcium response occurs, and *ρ* and *τ* are dimensionless variables for space and time, respectively. Additionally, the dependent variables are scaled in the following way:

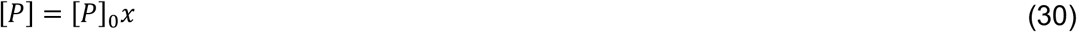

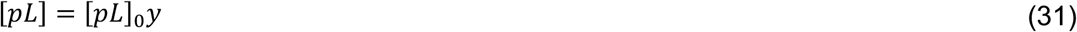

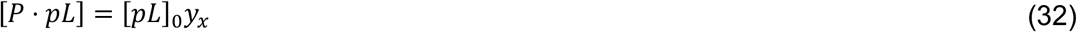

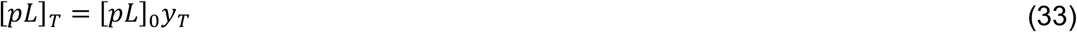

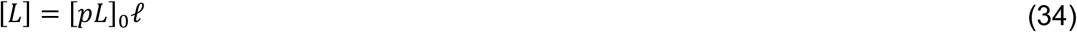

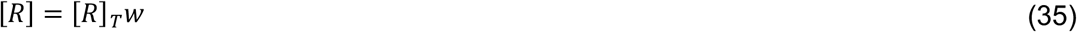

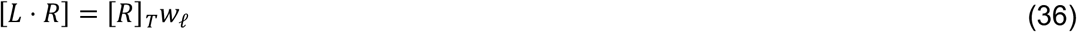

where 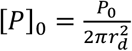, and *x,y,y_x_,y_T_,ℓ,w*, and *w_ℓ_* are dimensionless variables.

By applying these scales (28) – (36)to the model equations (16) – (22) one arrives at the following non-dimensionalized equations

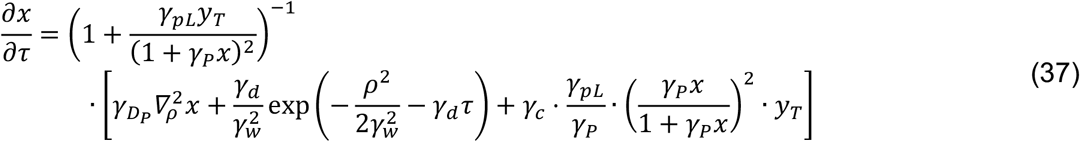

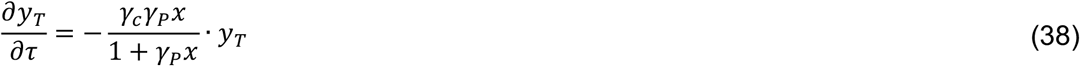

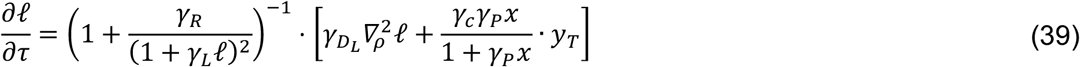

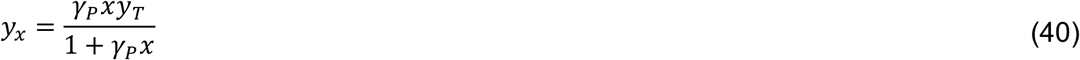

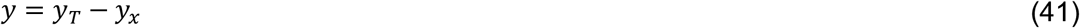

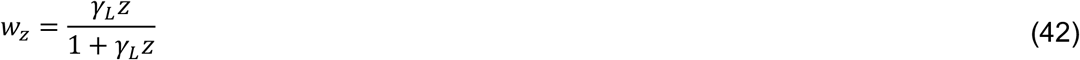

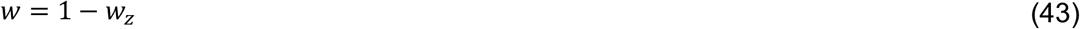

where we now have nine dimensionless parameters (see also Fig. S7, which highlights the influence of each of these parameters on model output)

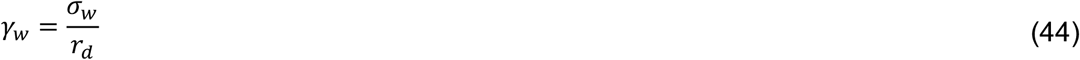

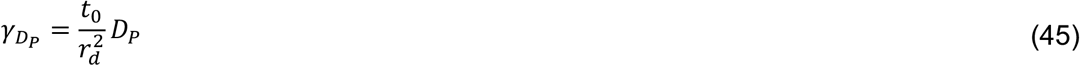

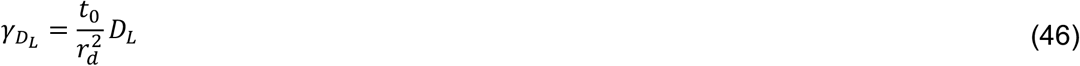

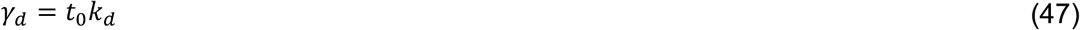

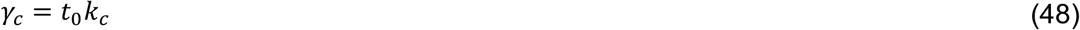

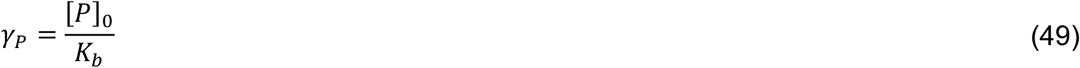

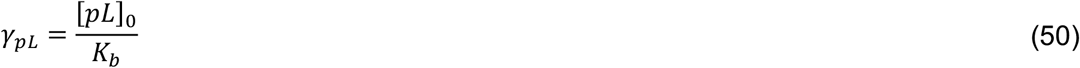

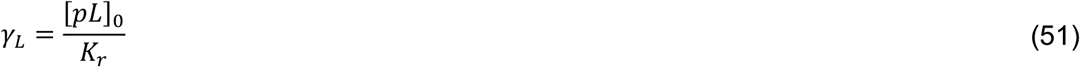

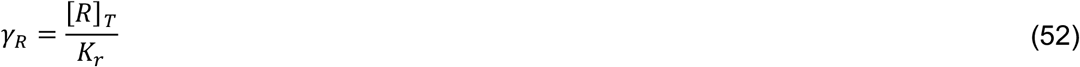

Additionally, the scaled initial and boundary conditions are

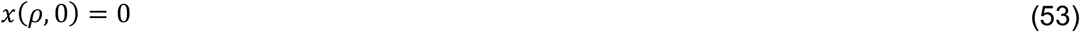

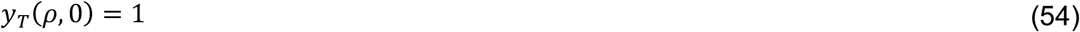

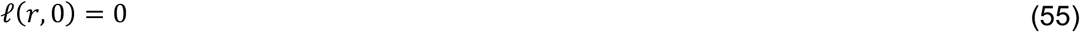

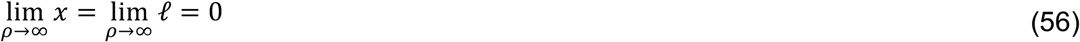

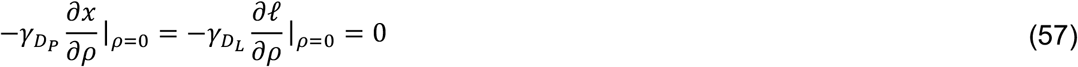

where, once again, initial conditions for the other scaled molecular components can be determined from the algebraic relations, although these are not important for solving the differential equations.

### Method of Solution

Due to the 1/*ρ* dependence of the Laplacian operator in polar coordinates, the boundary condition at *ρ* = 0 cannot be handled numerically in a simple fashion. Therefore, the boundary conditions were imposed at a small radial distance of *Δρ* = .001, and the protease source was shifted this distance as well so that the scaled source term becomes 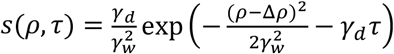. Additionally, the boundary condition at infinity must be imposed at a finite value *ρ_far_*. A value of *ρ_far_* = 100 (*r_far_* = 1000 *μm*) was chosen for simulations since values of *p_far_* larger than this do not cause large changes in the model output on the timescale of interest.

The software Mathematica was used to solve the above set of model equations. Since the system of equations is nonlinear and the equation for *y_T_* does not contain derivatives with respect to *ρ,* the method of lines was chosen to solve the set of differential equations. Because the spatial region is one-dimensional, the “Tensor Product Grid” method was used to discretize the spatial variable. The number of grid points for this discretization was chosen to be the smallest value so that use of a larger number of points did not yield a significantly different solution. To aid in any reproduction, the numerical differential equation solver function ‘NDSolveValue’ was used with settings specified by

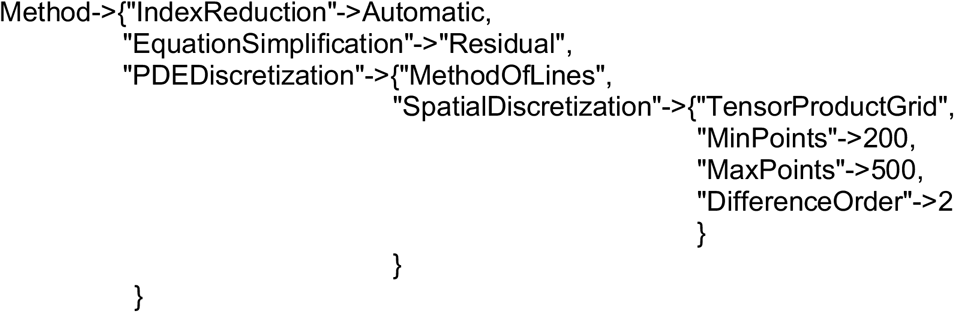

A more detailed description of the numeric solution algorithms and settings can be found in the Mathematica documentation.

### Model Output and Signal Radius

Solving the set of non-dimensionalized model equations along with a set of parameter values yields scaled molecular component concentrations as functions of *ρ* = *r/r_d_* and *τ* = *t/t*_0_. The model assumes that for a given spatial coordinate *ρ*, calcium will first be released into the cytosol when at least half of the receptors are bound to a ligand molecule. This sets a threshold for the scaled variable *w_ℓ_* as *w_thresh_* = 0.5. By solving for the radial distance *ρ_thresh_* such that *W_ℓ_*(*ρ_thresh_, τ*) = *W_thresh_*, we can then determine the signal radius as a function of time *ρ_thresh_*(*τ*). Therefore, it is assumed that *ρ_thresh_*(*τ*) is equal to the calcium radius during the distal calcium response.

After the distal calcium response, the calcium dynamics are determined by processes not covered in the current model. Therefore, it should be noted that the proposed model is not valid past the distal calcium response, as the ligand is not the sole factor in determining the calcium dynamics past this point.

### Estimate of Gbp diffusion constant

To estimate the free diffusion constant of the ligand Gbp, we assume that the ligand of molecular weight as 2.79 kDa ≈ 4.63 x 10^-21^ g (based on the peptide sequence described above) is spherical with a typical protein density of 1.37 g/cm^3^. This results in an estimated effective radius of 0.93 nm. Using the Stokes-Einstein relation at a temperature of 293 K in water results in an estimated diffusion constant of about 260 μm^2^/s, which is 122.2 in scaled units. This estimate is only used to approximate the Gbp diffusion constant; this parameter was still allowed to vary during the fitting process.

### Initial Guess Selection

Initial guesses for each selected distal calcium response was obtained by randomly searching parameter space for parameter sets that produced model outputs that met two criteria. First, the parameter set had to produce a distal calcium response that did not extend past the maximum of the distal calcium response to be fit to. Second, the parameter set had to produce a distal calcium response that had a sum of squares error that was 10% or less than the sum of squares error of a no-response model output (where the fraction of receptors bound to ligand never crosses the 50% threshold). Sum of square errors are determined according to step 2 of “Fitting the model to the data” below. For each of the four selected data sets to fit to, 32 initial guesses were obtained.

### Fitting the model to the data

The first goal of the model is to reproduce the distal calcium response data. This is done by fitting the model output of *_ρthresh_*(*τ*) to the data by varying the nine dimensionless parameters mentioned earlier. Due to a possibly large space of feasible parameter values, a fitting method is desired that searches a sufficient amount of the parameter space without taking too much time or biasing one region of parameter space over another. Therefore, the fitting method is based on orthogonal array sampling. The fitting steps are outlined below:

1. A distal calcium response dataset is chosen that the model will be fit to. In order to prevent *ρ_thresh_*(*τ*) from increasing to *ρ* values past the distal calcium response at later times, additional “constraint points” are appended to the dataset that are equal to the maximum extent of the distal calcium response. The number of constraint points is equal to the number of experimental data points in order to equally weigh fitting the distal calcium response with not extending past its maximum extent.
2. A fitting function to be minimized is defined in two parts. For the original data points, the fitting function is just a sum of squares difference between the data and the model. For the constraint points, the fitting function is also a sum of squares difference between the model and the data, but it is 0 if *ρ_thresh_*(*τ*) is less than the corresponding constraint point. This way there is only a penalty when the model output extends past the maximum extent of the distal calcium response. The goal of the fitting procedure is then to find a parameter set that minimizes the fitting function. More explicitly, we want to minimize the following piecewise function for data points (original and constraint) *d*(*τ*i**) and model output points *ρ_thresh_*(*τ_i_*) = *m*(*τ_i_*)

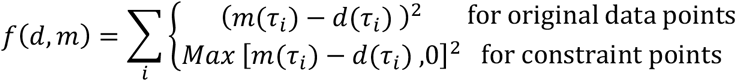
3. In order to vary the dimensionless parameters across desired ranges, some of which span orders of magnitude, the dimensionless parameters are replaced in the following way

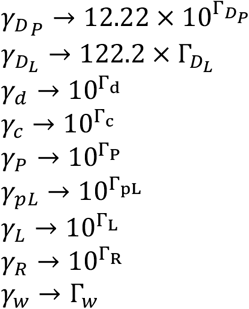

Therefore, we have created a new dimensionless parameter set Γ = {Γ_*D_P_*_, Γ_*D_L_*_, Γ_*d*_, Γ_*P*_, Γ_*P*_, Γ_*p_L_*_, Γ_*L*_, Γ_*L*_, Γ_*R*_, Γ_*W*_,} to use for the fitting method.
4. An initial set of parameter values Γ_0_ is determined as explained in the section “Initial Guess Selection” above.
5. An orthogonal array *A* is created with 729 runs, 9 factors, 3 levels, a strength of 5, and an index of 3. The levels are set to be {-1,0, 1}. i.e. *A* is a 729 x 9 matrix with the three entries −1,0, or 1 such that every 729 x 5 subarray of *A* contains every 5-tuple based on {-1,0,1} exactly 3 times.
6. For each row *A_i_* of the orthogonal array *A,* a new parameter set Γ_*i*_ is obtained by adding *A_i_* to the initial parameter set Γ_0_. Each of these parameter sets are then applied to the model, thus resulting in a set {*ρ_i_*(*τ*)} of 729 different *ρ_thresh_*(*τ*) functions.
7. For each *ρ_i_*(*τ*), the fitting function from step 2 is evaluated. The parameter set Γ_*i*_ that results in the best fit *ρ_i_*(*τ*) now becomes Γ_0_.
8. Steps 6 and 7 are repeated until the best fitting parameter set is Γ_0_
9. Steps 6 - 8 are repeated with entries {-1,0,1} in the orthogonal array replaced with entries {-1/2^*n*^, 0,1/2^*n*^} for *n* values 1,2, 3, and 4. The final set Γ_o_ is then determined to be the parameter set that best fits the data for the given initial parameter set.

### Distal calcium response properties from the model

The distal calcium response has two characteristics of interest: its time delay *t*_0,min_ and its effective diffusion constant *α_eff_*. It has been shown previously that the distal calcium response can be fit to an empirical “delayed diffusion equation” where *t*_0,min_ and *α_eff_* are independent parameters that are chosen to fit the data^8^. As shown inf Fig. 7F-F’, these measures appear to have a wound size dependency.

The parameters in the model that are affected by the wound size are the total amount of protease that is released and the spatial extent of the protease source. However, it is assumed that the protease per cell (area) should remain constant with respect to wound size. More specifically, in order to keep the protease density constant as the wound size changes, it must be that if *σ_w_* is scaled by some factor *λ,* then *P*_0_ must be scaled by *λ*^2^. These parameters appear in the dimensionless parameters 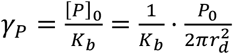 and 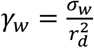. Therefore, scaling the wound size by a factor *λ* in the model corresponds to scaling *γ_w_* by *λ* and scaling *γ_P_* by *λ*^2^.

Starting with a best fit parameter set, the wound size was then varied using scaling factors of *λ* from 0.5 to 1.5 in step sizes of 0.1. An example of the calcium signal radius for each scaling factor is shown in Fig. 7E. Values of *t*_0,min_ and *α_eff_* were then determined for each new parameter set by fitting the delayed diffusion model to the RD model output. The delayed diffusion model, discussed in more detail in ^8^, describes the calcium signal radius by the equations

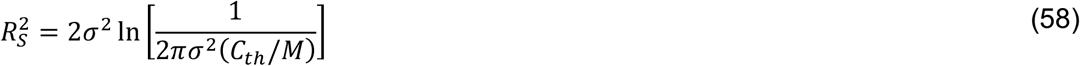

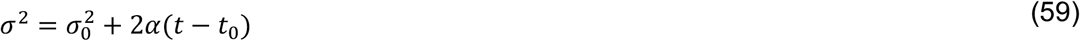

The effective diffusion constant *α* was obtained by directly fitting the RD model output to equations (58) and (59), while the time delay *t*_0,min_ was obtained from fits to (58) and (59) by

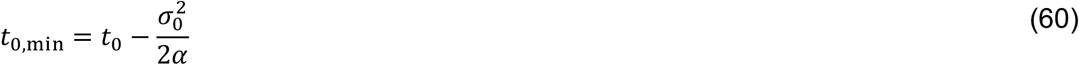

In other words, the time delay is the time when *σ* from equation (59) is equal to 0 given the best fit parameters to equations (58) and (59).

## Supplemental Movie Legends

**Movie S1: Calcium signaling is observed in response to wounds in *Drosophila* pupae.** Calcium response to wounds, monitored by GCaMP6m, is symmetric on both sides of the *pnr* boundary (yellow line) in the absence of gene knock-down *(pnr>mCherry-NLS).* The maximum radius of the rapid first expansion is marked by green circle. Scale bar = 50 μm.

**Movie S2: The distal calcium response requires components of the Gq-signaling pathway.**

Calcium response to wounds is monitored by GCaMP6m. The maximum radius of the rapid first expansion is marked by green circle. The distal calcium response only occurs in the control domain of each movie. Scale bar = 50 μm.

**Movie S3: Puncture wounds recapitulate laser wounds in *Drosophila* pupae.**

Calcium response to wounds is monitored by GCaMP6m in response to puncture by an electrolytically sharpened needle. Pupal nota were imaged with a 5X objective to give working distance for manual puncture. Scale bar = 100 μm.

**Movie S4: The distal calcium response can jump from the Gq knockdown domain to the control domain.**

Wound targeted within the *Gaq* knock-down domain (magenta, left of yellow line). Calcium response to wounds is monitored by GCaMP6m (green). The maximum radius of the rapid first expansion is marked by green circle. The distal calcium response is absent from the *G_aq_* knockdown domain until the signal diffuses into the nearby control domain (right of the yellow line). Scale bar = 50 μm.

**Movie S5: The distal calcium response requires the GPCR Methuselah-like 10.**

Calcium response to wounds is monitored by GCaMP6m. The maximum radius of the rapid first expansion is marked by green circle. The distal calcium response only occurs in the control domain. Scale bar = 50 μm.

**Movie S6: Gbp1 and Gbp2 elicit a calcium response in wing discs in an Mthl10-dependent manner.**

Control (white outline) and *mthl10* knockdown (red outline) wing discs mounted together in the same media bubble. Gbp1 added at t = 0 seconds to 5 nM final concentration (left), or Gbp2 added at t = 0 seconds to 50 nM final concentration (right), elicits a calcium response in only the control disc, monitored by GCaMP6m. Scale bar = 100 μm.

**Movie S7: *Drosophila* extract elicits a calcium response in wing discs in an Mthl10-dependent manner.**

Control (white outline) and *mthl10* knockdown (red outline) wing discs mounted together in the same media bubble. Adult fly extract (left) or larval extract (right), added at t=0 to 5% final concentration, elicits a calcium response in only the control discs, monitored by GCaMP6m. Scale bar = 100 μm.

**Movie S8: Larval extract elicits a calcium response in wing discs in a Gbp-dependent manner.**

Control (white outline) and *ΔGbp1,2* (red outline) wing discs mounted together in the same media bubble. *ΔGbp1,2* larval extract, added at t=0 to 5% final concentration, elicits a calcium response in only the control disc, monitored by GCaMP6m. Scale bar = 100 μm.

**Movie S9: Model output of scaled molecular component concentrations and calcium signal radius vs. time**

(A) Protease concentration scaled by the concentration of total protease within an area of ~638 μm^2^ (see supplementary information). (B) Free pro-ligand and ligand concentrations scaled by the initial pro-ligand concentration. (C) Free receptor and bound receptor concentrations scaled by the total receptor concentration. The bound receptor threshold for triggering calcium release is taken as 0.5 (dashed line). Red point indicates where the bound receptor concentration crosses this threshold. (D) Calcium signal radius as a function of time as determined from the point in space where the bound receptor crosses the signaling threshold. Red point indicates the signal radius at the corresponding point in time.

## Notes

### Competing Interest Statement

The authors have declared no competing interest.

